# Host Phosphatase PPM1G Governs Influenza vRNP Homeostasis by Orchestrating NP Phosphorylation Dynamics and Autophagy

**DOI:** 10.1101/2025.09.18.676991

**Authors:** Mengmeng Yu, Yuxing Qu, LiuKe Sun, Zhenyu Zhang, Rongkuan Sun, Xing Guo, Huiling Ren, Xiaojun Wang

## Abstract

The influenza viral ribonucleoprotein (vRNP) serves as the machinery for viral RNA replication. It self - assembles from the RNA - dependent RNA polymerase (RdRp), viral RNA, and nucleoprotein (NP). During vRNP replication, NP proteins must rapidly switch between monomeric and polymeric states to sustain dynamic equilibrium, which relies on NP phosphorylation-dephosphorylation cycling and is crucial for viral nucleic acid replication. However, the regulatory mechanisms controlling this process remain poorly understood. In this study, we employed an RdRp/vRNP-targeted affinity mass spectrometry-based differential analysis method, discovered that the cellular protein phosphatase Mg^2+^/Mn^2+^-dependent 1G (PPM1G) is a phosphatase of NP and required for maintaining viral polymerase activity. Overexpression of PPM1G decreased viral replication. In *ppm1g* ^flox/flox^-Sftpc Cre mice, lethal influenza infection led to survival and mild disease, as well as reduced lung viral loads and inflammation. These results suggest that PPM1G is a key host molecule for maintaining steady state of viral replication. PPM1G expression is upregulated upon influenza virus infection. This heightened expression occurs in response to increased viral protein production, simultaneously leading to elevated dephosphorylation of NP, which accelerates NP polymerization to promote viral replication. Meanwhile, surplus NP can be degraded through the ATG7 autophagylysosome pathway. Our research elucidates a mechanism that PPM1G maintains viral replication homeostasis by regulating NP status and fate.

## Introduction

Influenza A virus (IAV) and influenza B virus (IBV) are the main pathogens causing human seasonal influenza, posing a huge threat to global public health and economic development[1–5]. The viral ribonucleoprotein complex (vRNP) serves as the minimal functional unit for influenza virus replication, comprising the viral RNA-dependent RNA polymerase (RdRp), nucleoprotein (NP), and viral RNA (vRNA). The RdRp consists of three heterotrimeric proteins: PB1, PB2, and PA[6–10]. The viral RNA-dependent RNA polymerase (RdRp) enables viral genome replication through a mechanism mediated by acidic nuclear phosphoprotein 32A (ANP32A), whereby ANP32A facilitates the formation of an asymmetric RdRp dimer. In this structure, one polymerase subunit is dedicated to RNA synthesis by binding the vRNA template, while the partner subunit captures the newly synthesized RNA and initiates its assembly with NP for vRNP formation [11]. vRNP assembly is essential for influenza virus replication, transcription, infection, and adaptability, forming a pivotal step in its life cycle.NP is an important cofactor that binds with the vRNA to form vRNP [6, 12].

NP undergoes oligomerization to assemble a double-helical framework, with the central region of vRNA is encapsidated by NP. This NP-vRNA assembly forms the structural core of the vRNP, serving as a stable platform to support replication and transcription activities mediated by RdRp, with the polymerase anchored at one terminus of the vRNP [6, 13]. During replication initiation, a subset of NP molecules transiently dissociates from the vRNA, enabling RdRp to directly access the vRNA template and initiate complementary RNA (cRNA) synthesis. RdRp utilizes the cRNA as a template to generate negative strand vRNA, while NP concomitantly associates with the nascent vRNA, ultimately forming progeny vRNP complexes [12, 14–16]. NP alternates between tightly and loosely bound states during replication stages, with this structural dynamism governing viral replication/transcription efficiency. The monomer-oligomer transition of NP, a core mechanism underpinning vRNP functionality, maintains a dynamic equilibrium of dissociation-rebinding interactions[17–20]. During vRNP assembly, the NP tail loop conformation is changed to promote interaction with another NP binding groove, forming multiple oligomerized NPs. vRNA wraps around the polymerized NPs and assembles with RdRp to exert replication function[19, 21–24] .

The dynamic transition between phosphorylation and dephosphorylation of NP is key to its regulation of monomer and oligomerization states[17, 18, 25]. The phosphorylated form of NP tends to adopt a monomeric state, and the formation of this monomer is regulated by the host protein kinases, with PKCδ—a member of the protein kinase C (PKC) kinase family—being recruited by PB2 to NP’s tail loop. This PB2-mediated recruitment of PKCδ augments IAV-NP phosphorylation, thereby suppressing NP oligomerization and ultimately impeding vRNP assembly[25]. The oligomerization form of NP predominantly exists in a dephosphorylated state. In this conformation, the NP oligomerization becomes entwined with vRNA, forming a stable vRNP that effectively shields the viral genetic material from degradation by host nucleases. Dephosphorylated NP is a prerequisite for oligomerization, however, the key regulatory enzymes to dephosphorylate NP and the mechanisms in NP cycling remain largely unknown.

Utilizing an IBV RdRp/vRNP affinity purification mass spectrometry (AP-MS)-based differential analysis approach, which quantifies abundance discrepancies of identical proteins across distinct vRNP complexes, we identified significant enrichment of PPM1G, a member of the PPM (Protein Phosphatase Mg^2+^/Mn^2+^-dependent) family[26], in IBV-associated vRNP. Both overexpression and knockout of PPM1G at the cellular level significantly reduce influenza viral polymerase activity and replication, indicating its dual roles as a critical regulator of vRNP functional homeostasis. In vivo experiments in mice have also demonstrated that PPM1G is required for influenza virus replication. Genetic ablation of PPM1G in mice (*ppm1g ^flox/flox^-*Sftpc Cre mice; flox: loxP-flanked; Sftpc: Surfactant Protein C; cre: Cre recombinase) conferred resistance to lethal influenza challenge, demonstrating markedly improved survival rates compared to *ppm1g* ^flox/flox^. Mechanistically, PPM1G contains a unique acidic amino acid repeat motif that distinguishes it from other PPM family proteins, and this motif may enable its direct interaction with vRNP or viral polymerase subunits (PB1, PB2) and NP to exert its functional role. The phosphatase activity of PPM1G is indispensable for orchestrating influenza replication by dephosphorylating NP to facilitate its oligomerization—a prerequisite for vRNP assembly and subsequent genome packaging. Furthermore, PPM1G mediates NP turnover through an ATG7-dependent autophagosome-lysosome pathway, suggesting this post-translational modification circuit may represent a cycling mechanism for NP. Our findings reveal that PPM1G expression modulates the state and fate of NP, thereby maintaining the functional homeostasis of vRNP complexes during influenza virus replication.

## Results

### 1. PPM1G plays an important role in maintaining virus growth in vitro

To identify key cellular factors that **are** involved in the vRNA working status, three groups of plasmids, Group a (RdRp with Flag-tagged), Group b (vRNP with Flag-tagged), and Group c (vRNP without tagged), were transfected into HEK293T cells, respectively. Flag-tagged magnetic beads were used for co-immunoprecipitation (Co-IP) to purify the RdRp and vRNP complexes. Following mass spectrometry analysis, the protein abundance of each component is shown in Figure 1A, and significant variations in protein abundance were identified among Group a, b and c. Using mass spectrometry analysis, we identified 15 peptide segments of the IBV-NP protein in Group b (Figure 1B). The NP protein exhibited higher abundance in Group b and was absent in Group a. Concurrently, we observed that the core proteins of the RdRp complex—PA, PB1, and PB2—were present in both Groups a and b but were differentially expressed between the two groups (Figure 1C), further confirming the specific association of Group b with vRNP. Designating Group a as the reference control, we conducted comparative analysis focusing on Group b to identify host proteins specifically or differentially associated with vRNP relative to Group a. This screening revealed PA, PB2, PB1, NP, ANP32B, ANP32A, PPM1G, SET, STOML2, ANP32E, SSRP1, and RPS27A (Figures 1C, D), with their respective protein abundance profiles illustrated in Figure 1C. Beyond the viral proteins PA, PB1, PB2, and NP, vRNP-specific binding proteins were identified, including members of the ANP32 family (ANP32A, ANP32B, ANP32E), ANP32 family proteins play a decisive role in the process of influenza virus replication[27–33], suggested that the mass spectrometry results were highly reliable. In addition to the ANP32 family proteins, we also identified four novel host proteins with extremely high abundance as candidate proteins of interest in this study: PPM1G, SET, STOML2 and SSRP1(Figure 1D). Additionally, we observed that when using Group c as the background control group, these proteins in Group a (Figure S1A) and Group b (Figure S1B) exhibited significantly higher abundance compared to Group c. This further supports the reliability of these candidate proteins.

**Figure 1:**
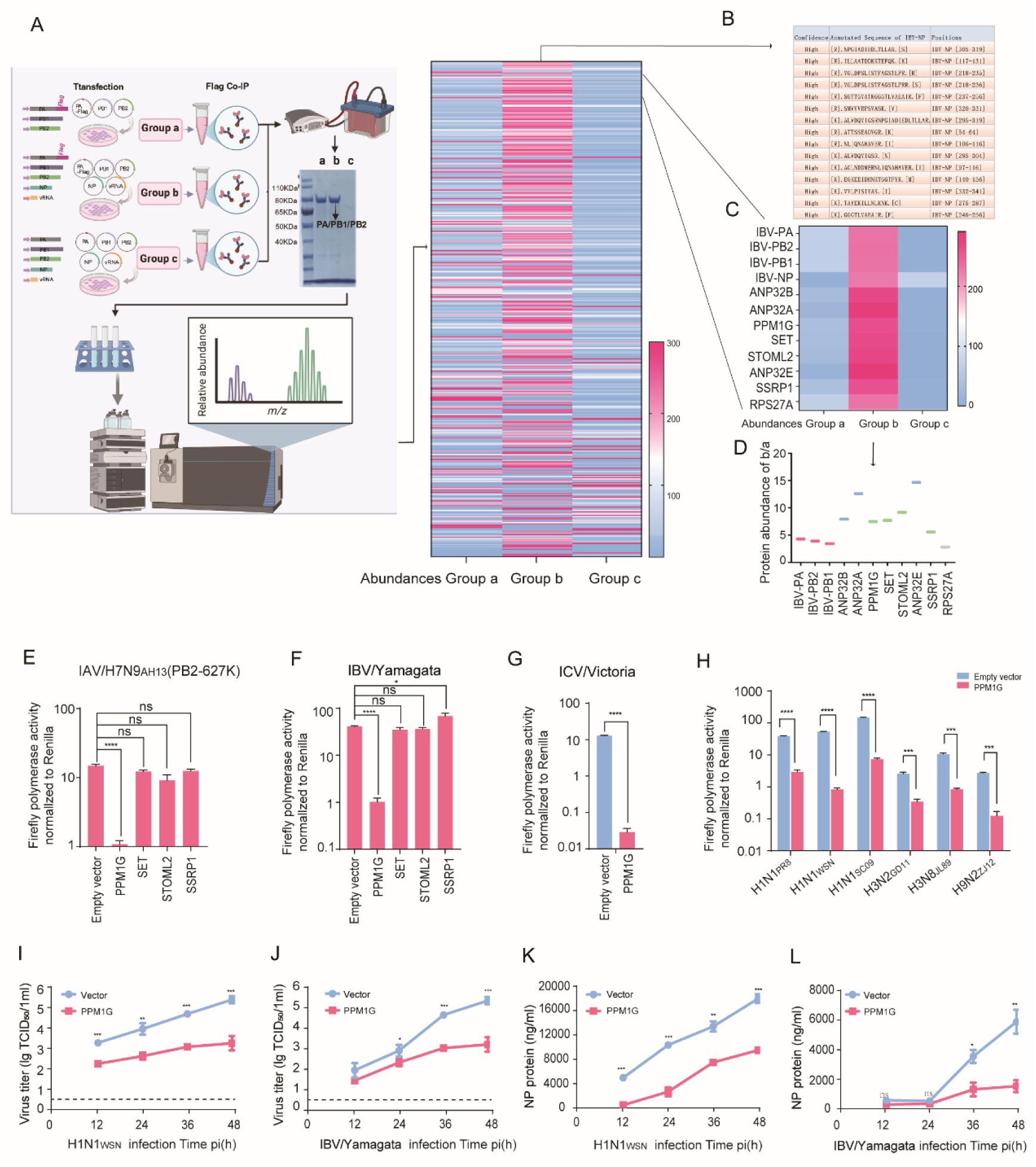
PPM1G is associated with vRNP and can significantly inhibit influenza virus polymerase activity and replication when overexpressed. A-D : Following transfection of HEK293T cells with Flag-tagged RdRp (Group a), Flag-tagged vRNP (Group b) or untagged vRNP (control, Group c), co-immunoprecipitation (Co-IP) was performed using Flag magnetic beads. The immunoprecipitated samples were then resolved by SDS- PAGE and stained with Coomassie brilliant blue staining. Proteins with differential expressions across groups a, b, and c were identified by mass spectrometry (MS) analysis (A). In Group b, MS identified 15 peptide fragments of IBV-NP (B). Differential proteomic analysis between RdRp and vRNP identified several associated proteins including PA, PB2, PB1, NP, ANP32B, ANP32A, PPM1G, SET, STOML2, ANP32E, SSRP1 and RPS27A (D), whose relative abundance is presented in C. E-H: Overexpression of PPM1G protein significantly inhibits the activity of polymerases from influenza A, B and C viruses. Minigenome assays of H7N9_AH13_(PB2-627K) (E), IBV-Yamagata (F), ICV/Victoria (G) and other subtypes of IAV (H) in human HEK293T cells. These cells were cotransfected with either empty vector or different Flag-tagged constructs. Error bars represent means ± SD from n = 3 independent biological replicates; Statistical significance was determined by two-tailed unpaired t-test. NS = not significant, \**p*<0.05, *** *p* < 0.001, **** *p* < 0.0001. I-L: Impact of PPM1G overexpression on IAV or IBV replication. HEK293T cells overexpressing PPM1G or empty control were infected with IAV (H1N1_WSN_) (MOI = 0.001) or IBV/Yamagata (MOI = 0.01). The viral titer (I, J) and the NP content (K, L) were measured using TCID_50_ and AcELISA. Error bars represent means ± SD from n = 3 independent biological replicates; Statistical significance was determined by Two-tailed unpaired t-test. NS = not significant, \**p*<0.05, ** *p* < 0.01, *** *p* < 0.001.

We found that overexpression of PPM1G, but not other proteins, significantly suppressed the polymerase activity in IAV/H7N9_AH13_ (PB2-627K), IBV/Yamagata and ICV/Victoria influenza viruses (Figures 1E–G). Moreover, PPM1G overexpression significantly inhibited polymerase activity across multiple subtypes of influenza A virus, including H1N1_PR8_, H1N1_WSN_, H1N1_SC09_, H3N2_GD11_, H3N8_JL89_ and H9N2_ZJ12_ (PB2-627K) (Figure 1H). Furthermore, PPM1G overexpression inhibited influenza virus polymerase activity in IAV/H7N9_AH13_ (PB2-627K), IBV/Yamagata and ICV/Victoria in a dose-dependent manner (Figure S2A–C). The polymerase activities of IAV/H7N9_AH13_ (PB2-627K), IBV/Yamagata and ICV/Victoria were also significantly reduced in PPM1G-stably expressing cell lines (in which the physiological state was consistent with wild-type HEK293T) (Figures S2D–F), consistent with transient transfection results. These findings demonstrate that PPM1G overexpression significantly inhibits the polymerase activity of influenza A, B, and C viruses. Additionally, PPM1G overexpression significantly suppressed viral replication (Figures 1I, J) and virus production (Figures 1K, L) in both H1N1_WSN_ and IBV/Yamagata viruses.

sgRNAs were designed for CRISPR-Cas9 knockdown targeting the *ppm1g* gene, and the *ppm1g* knockout cell line (*ppm1g*-KO cells) was successfully established (Figure 2A, B). Cell proliferation and physiological status in this cell line were consistent with those of the wild-type (Figure 2C).P*pm1g* knockout significantly reduced the activities of the polymerases from IAV/H7N9_AH13_ (PB2-627K), IBV/Yamagata and ICV/Victoria influenza viruses (Figure 2 D-F) as well as in *ppm1g* stable knockdown cell lines (*ppm1g*-sh cells) (Figure S2G, H). In *ppm1g*-KO cells, reconstitution of PPM1G restored full IBV/Yamagata polymerase activity, however, high level PPM1G overexpression again led to polymerase inhibition (Figure 2G). Reconstitution of PPM1G in KO cells restored viral infectivity and virus production in both IAV H1N1_WSN_ (Figure 2H, J) and IBV/Yamagata (Figure 2 I, K). These results suggest that *ppm1g* knockout/downregulation significantly inhibits influenza virus polymerase activity.

**Figure 2.**
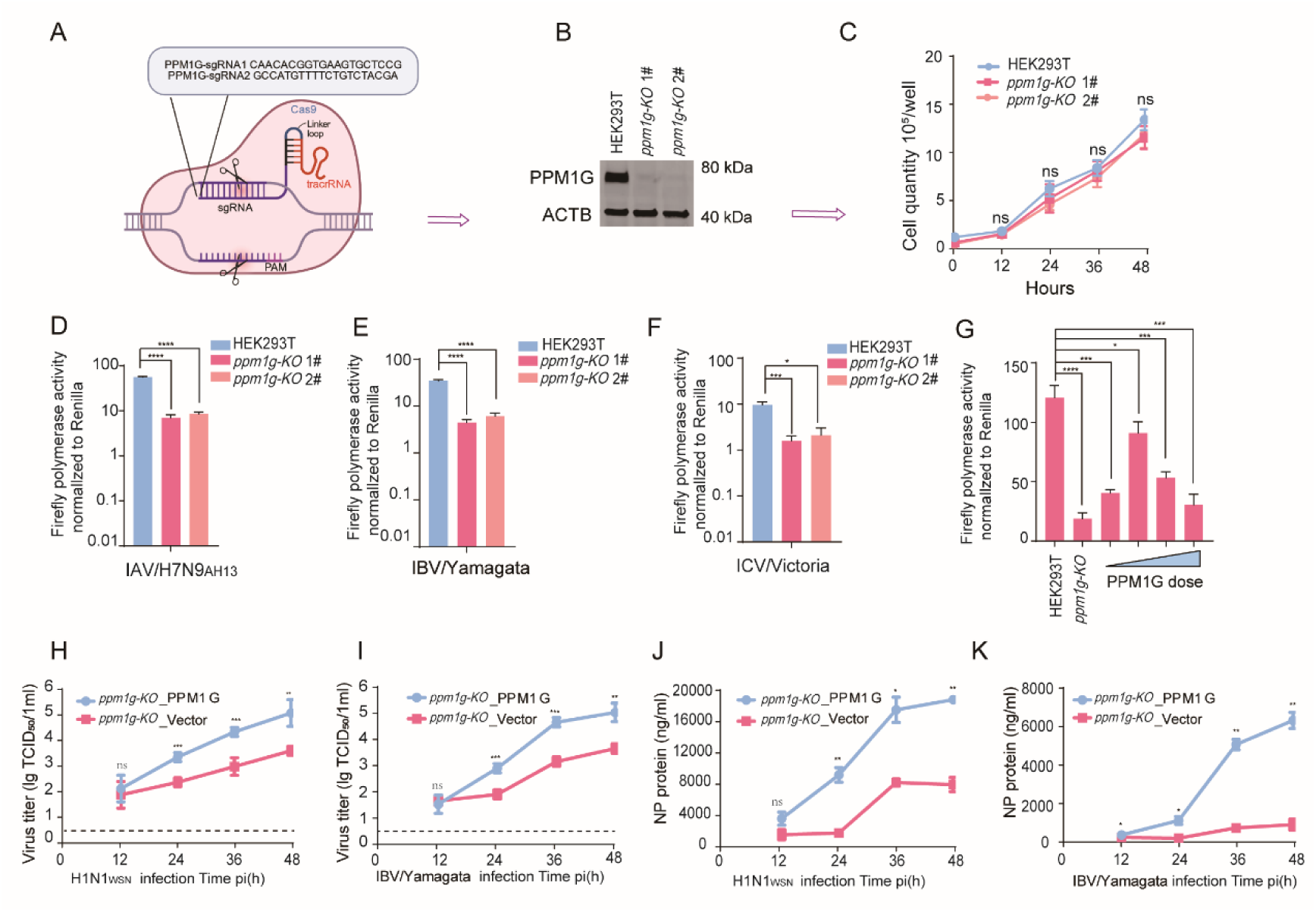
Knockout of *ppm1g* significantly inhibits influenza virus replication. A-G: *ppm1g* gene knockout significantly inhibits the activity of polymerases from influenza A, B and C viruses. Schematic diagram of sgRNA design for *ppm1g* (A). Detection of PPM1G protein expression in WT and *ppm1g*-KO cells (B), and detection of their physiological status (C). Minigenome assays of H7N9_AH13_(PB2-627K) (D), IBV-Yamagata (E) and ICV/Victoria (F) in WT and *ppm1g*-KO cell lines. Detection of IBV/Yamagata polymerase activity after PPM1G supplementation in *ppm1g*-KO cells (G). Error bars represent means ± SD from n = 3 independent biological replicates; Statistical significance was determined by two-tailed unpaired t-test. NS = not significant, \**p*<0.05, *** *p* < 0.001, **** *p* < 0.0001. H-K: Impact of *ppm1g* knockout on IAV or IBV replication. *ppm1g*-KO cells overexpressing PPM1G or empty control were infected with IAV (H1N1_WSN_) (MOI = 0.001) or IBV/Yamagata (MOI = 0.01). The viral titer (H, I) and the NP content (J, K) were measured using TCID_50_ and AcELISA. Error bars represent means ± SD from n = 3 independent biological replicates; Statistical significance was determined by Two-tailed unpaired t-test. NS = not significant, \**p*<0.05, ** *p* < 0.01, *** *p* < 0.001.

The above findings establish that influenza virus replication strictly depends on maintaining a proper level of PPM1G protein expression.

### 2. PPM1G promoted IAV and IBV replication in vivo

To evaluate the physiological relevance of PPM1G in vivo in combating influenza virus, a conditional knockout mouse model was used, as homozygous *ppm1g*−/− is likely to cause perinatal death (as indicated in the MGI (Mouse Genome Informatics) database). A lung expressing Cre line, surfactant associated with protein C-Cre (Sftpc-Cre), was crossed with *ppm1g* ^flox/flox^ mice to produce *ppm1g* ^flox/flox^-Sftpc Cre mice (Figure S3 A, B), which were genotypically confirmed by PCR (Figure S3 C-E) and WB (Figure S3 F, G).

In wild type mice infected with the WSN influenza virus, the protein expression level of PPM1G in the lungs was highly upregulated at 3 days post-infection along with high level expression of viral proteins (Figure 3A), suggesting that PPM1G may be required for viral replication. *Ppm1g* ^flox/flox^-Sftpc Cre and control *ppm1g* ^flox/flox^ mice were intranasally infected with 10 MLD_50_ of H1N1 _WSN_ virus, and their survival and body weight changes were monitored daily for 14 days (Figure 3B). The results showed that all 12 *ppm1g* ^flox/flox^ mice died by day 12. In contrast, 5 of 12 *ppm1g* ^flox/flox^-Sftpc Cre mice survived on day 14 (Figure 3C). Moreover, the body weight of *ppm1g* ^flox/flox^ mice continued to decline over 14 days until all died, while *ppm1g* ^flox/flox^-Sftpc Cre mice showed a sustained downward trend in body weight within 9 days after viral infection, however, starting from the day 10, the body weight of the mice gradually increased (Figure 3D). At 4 days post-infection, viruses from the lung tissues of *ppm1g* ^flox/flox^ mice and *ppm1g* ^flox/flox^-Sftpc Cre mice were extracted for viral growth curve analysis, RT-PCR, immunohistochemistry (IHC), and H&E staining. The results showed that the progeny virus yields from *ppm1g* ^flox/flox^-Sftpc Cre mice were significantly lower than those in the control group (Figure 3E), and the mRNA of the viral NP and M (matrix protein) was correspondingly reduced in *ppm1g* ^flox/flox^-Sftpc Cre mice (Figure 3F).

**Figure 3:**
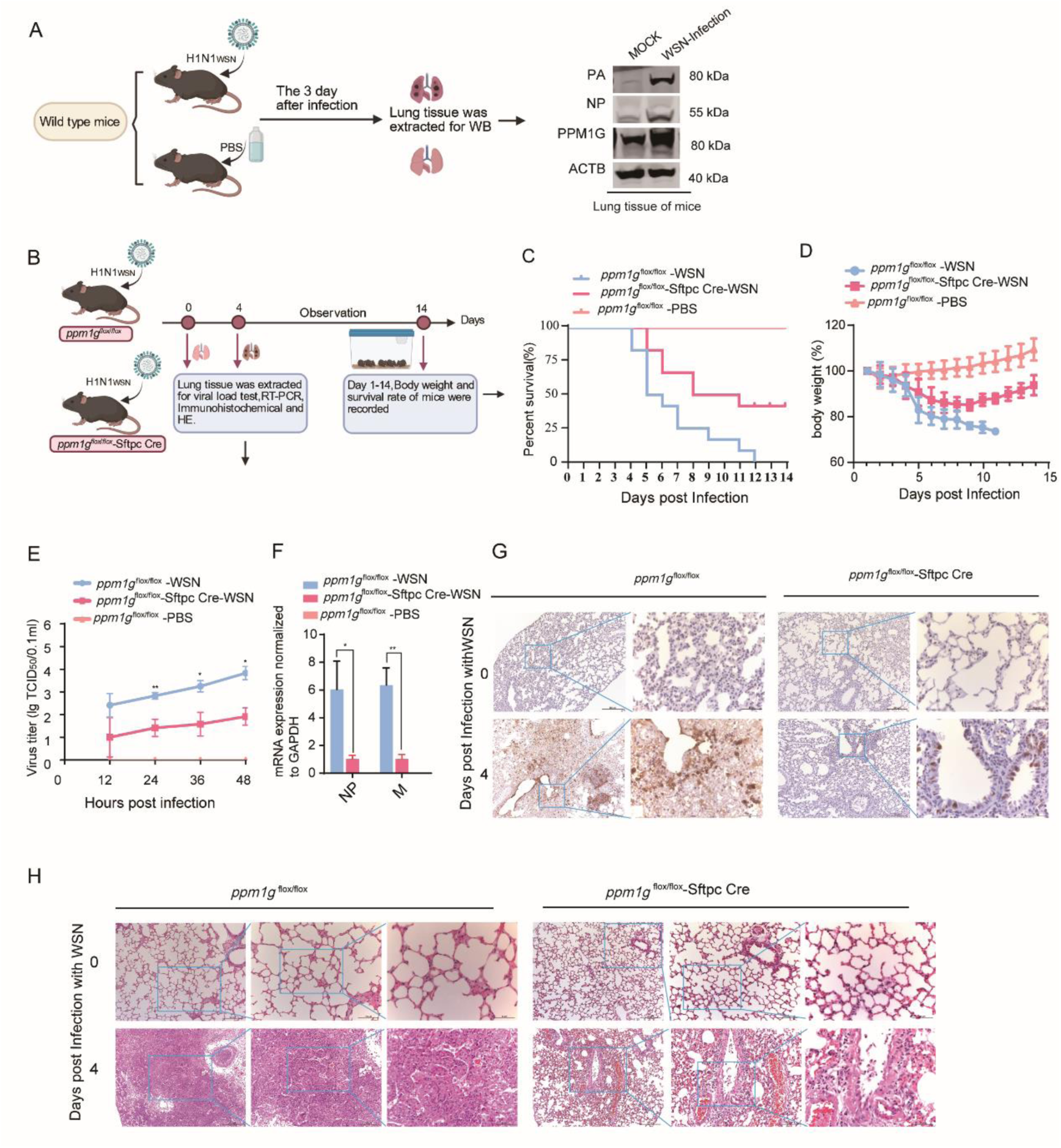
*Ppm1g*-deficient mice are resistant to influenza A virus infection. A: Three days after IAV infection of mice, western blot detection of viral and PPM1G protein expression in mouse lungs. B–D: (B) *ppm1g* ^flox/flox^-Sftpc cre and *ppm1g* ^flox/flox^ mice (5-6 weeks) were intranasally infected with IAV/H1N1_WSN_ to establish severe disease (10 MLD_50_; n = 12) and monitored for (C) survival and (D) body weight loss (mean ± SEM). Data presented were pooled from three independent experiments. Kaplan–Meier survival curves subjected to log-rank (Mantel–Cox)analysis. E: Viral titer in lung tissues from WSN-virus-infected *ppm1g* ^flox/flox^-Sftpc cre and *ppm1g* ^flox/flox^ mice at 4 dpi measured by TCID_50_ assay. Data are from three mice per group and are representative of three independent experiments, with each sample run in triplicate. F: Viral NP and M mRNA in lung tissues of WSN virus-infected *ppm1g* ^flox/flox^-Sftpc cre and *ppm1g* ^flox/flox^ mice at 4 dpi as determined by RT-qPCR. Data are presented as means ± SD and are representative of three independent experiments (n = 3 mice per group). G-H: H&E staining (G) and NP immunohistochemistry(H) of lung tissues from *ppm1g* ^flox/flox^-Sftpc cre and *ppm1g* ^flox/flox^ mice infected with WSN virus at 4 dpi; scale bars = 50 or 200 μm. Representative images from 3 mice per group from three independent experiments are shown.

Compared with *ppm1g* ^flox/flox^ mice, histopathological examination also showed that *ppm1g* ^flox/flox^- Sftpc Cre mice had less NP protein detected in the lungs, and the inflammatory response was alleviated (Figure 3G, H).

In an IBV infection model, IBV/Yamagata was adapted to mice to generate a mouse-adapted strain (IBV/MD). 10MLD_50_ of IBV/MD influenza virus was used to infect *ppm1g* ^flox/flox^ -Sftpc Cre and *ppm1g* ^flox/flox^ mice (Figure S4 A). 10 *ppm1g* ^flox/flox^ mice all died by day 5 after infection with sharp weight loss, while the 3 *ppm1g* ^flox/flox^-Sftpc Cre mice survived to day 14, and their weights began to recover from day 9 (Figure S4 B, C). The progeny virus yields and lung inflammatory infiltration in the *ppm1g* ^flox/flox^-Sftpc Cre mice were lower compared to the *ppm1g* ^flox/flox^ mice (Figure S4 D, E). Histopathological analysis revealed that compared to ppm1g ^flox/flox^ controls, ppm1g^flox/flox^-Sftpc Cre mice exhibited reduced NP protein detection in lung tissues along with attenuated inflammatory responses (Figure S4 F, G). In summary, PPM1G deficiency reduces IAV and IBV replication and alleviates the severity of lung infection in mice.

### 3. PPM1G binds to the viral polymerase and regulates polymerase activity through its acidic domain

The host protein PPM1G belongs to the PPM family. In this study, phosphatase proteins known from previous studies (PPM1A, PPM1B, PPM1D, PPM1J, PPM1K and PPM1L) were selected for further investigation (Figure 4A). We found that, compared with other PPM family proteins, the overexpression of PPM1G protein significantly down-regulated the activity of polymerases of influenza A, B and C viruses (about 20-100 times), although overexpression of PPM1A, PPM1B, and PPM1J each slightly decreased influenza virus polymerase activity (Figure 4B-D). Subsequently, we found that only PPM1G knockdown in A549 cells significantly inhibited the polymerase activity of influenza A, B, and C viruses, whereas PPM1A, PPM1B, and PPM1J did not (Figure 4E-H). This further confirmed that among the PPM family, PPM1G is uniquely capable of regulating influenza viral polymerase activity.

**Figure 4:**
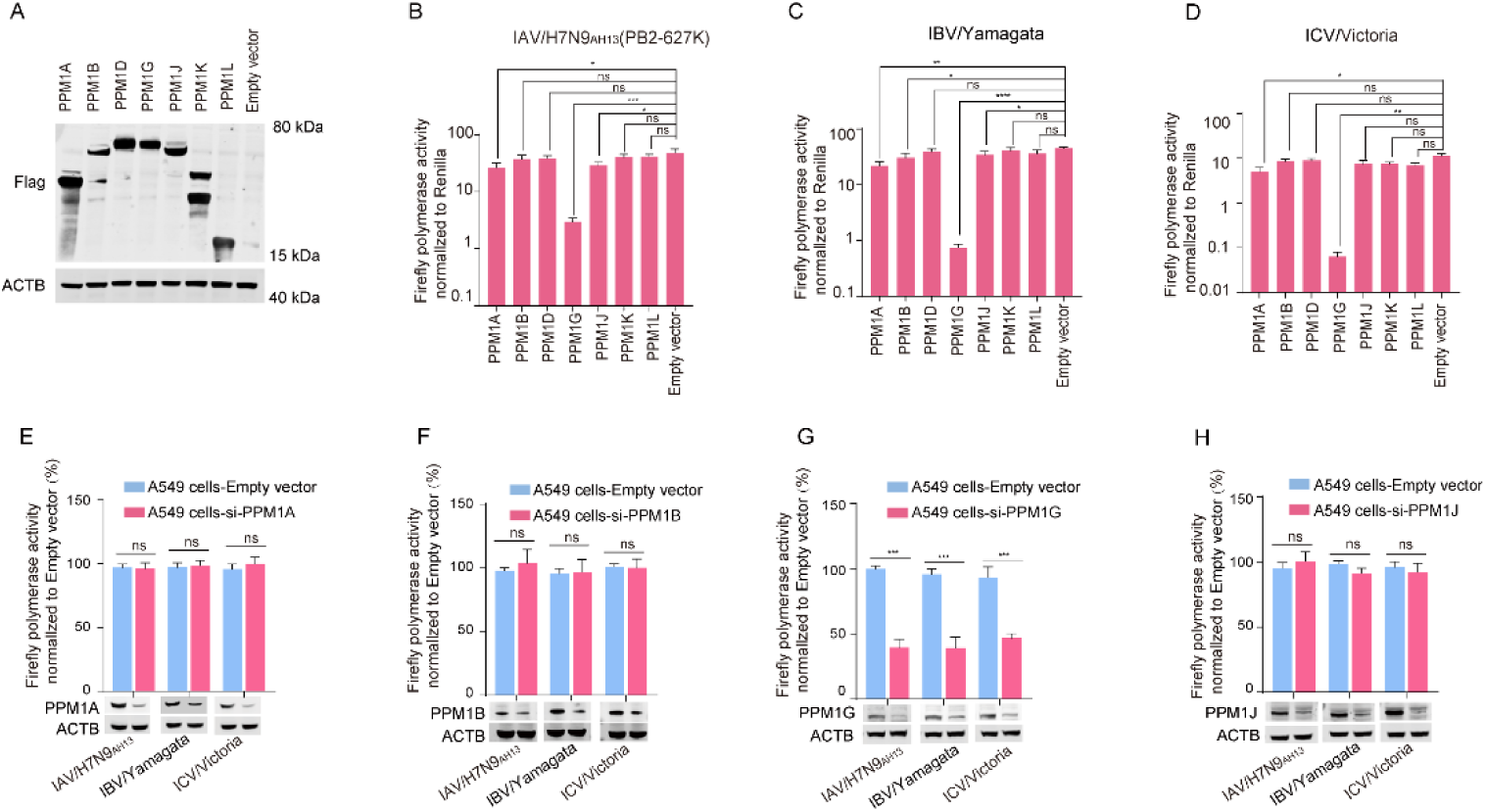
Effect of overexpression of PPM family proteins on influenza virus polymerase activity. A: Detection of PPM1A, PPM1B, PPM1D, PPM1G, PPM1J, PPM1K and PPM1L protein expression using WB. B-D: Minigenome assays of activities of polymerases from H7N9_AH13_(PB2-627K) (B), IBV-Yamagata (C) and ICV/Victoria (D) in human HEK293T cells co-expressing different Flag tagged constructs of PPM family proteins. Error bars represent means ± SD from n = 3 independent biological replicates; statistical significance was determined by two-tailed unpaired t-test. NS = not significant, \**p*<0.05, ** *p* < 0.01, *** *p* < 0.001, **** *p* < 0.0001. E-P: Detection of the effect of siRNA targeting PPM1A (*siPPM1A*) (E), siRNA targeting PPM1B (*siPPM1B*) (F), siRNA targeting PPM1G (*siPPM1G*) (G), siRNA targeting PPM1J (*siPPM1J*) (H) on the polymerase activity of H7N9AH13(PB2-627K), IBV-Yamagata and ICV/Victoria in A549 cells. Error bars represent means ± SD from n = 3 independent biological replicates; statistical significance was determined by two-tailed unpaired t-test. NS = not significant, *** *p* < 0.001.

Our previous work demonstrated that PPM1G significantly regulates the polymerase activity of IAV, IBV, and ICV. Subsequently, we used Co-IP to investigate which protein within the vRNP interacts with PPM1G. The results showed that IBV-NP, PB1, and PB2 could bind to PPM1G (Figure 5A-C). PPM1G also interacted with the IBV vRNP or with the PA+PB1/PB1+PB2 subunits (Figure S5A-B). To understand why PPM1G, compared to other family proteins, specifically regulates influenza virus replication, we used online analysis tools (https://www.uniprot.org/ and https://swissmodel.expasy.org/interactive) to analyze the domains and 3D structures of PPM family proteins. We found that PPM1G possessed a unique acidic amino acid repeat sequence at positions 258-322 (rich in DE [Asp/Glu]), which was not found in the other PPM family proteins [34] (Figure 5D,E). We hypothesized that this domain might be a key region for PPM1G’s regulation of influenza viruses. As expected, PPM1G-mut (lacking amino acids 258-322) failed to interact with NP, PB1, or PB2(Figure 5F, G). We then investigated the effect of PPM1G-mut on polymerase activity. Compared to wild-type PPM1G, the mutant PPM1G-mut lost its ability to regulate the polymerase activity of both IAV/H7N9_AH13_ (PB2-627K), IBV/Yamagata, and ICV/Victoria (Figure 5H-J). All these results indicated that the unique acidic amino acid repeat domain (positions 258–322) of PPM1G is crucial for its function in regulating influenza virus replication

**Figure 5:**
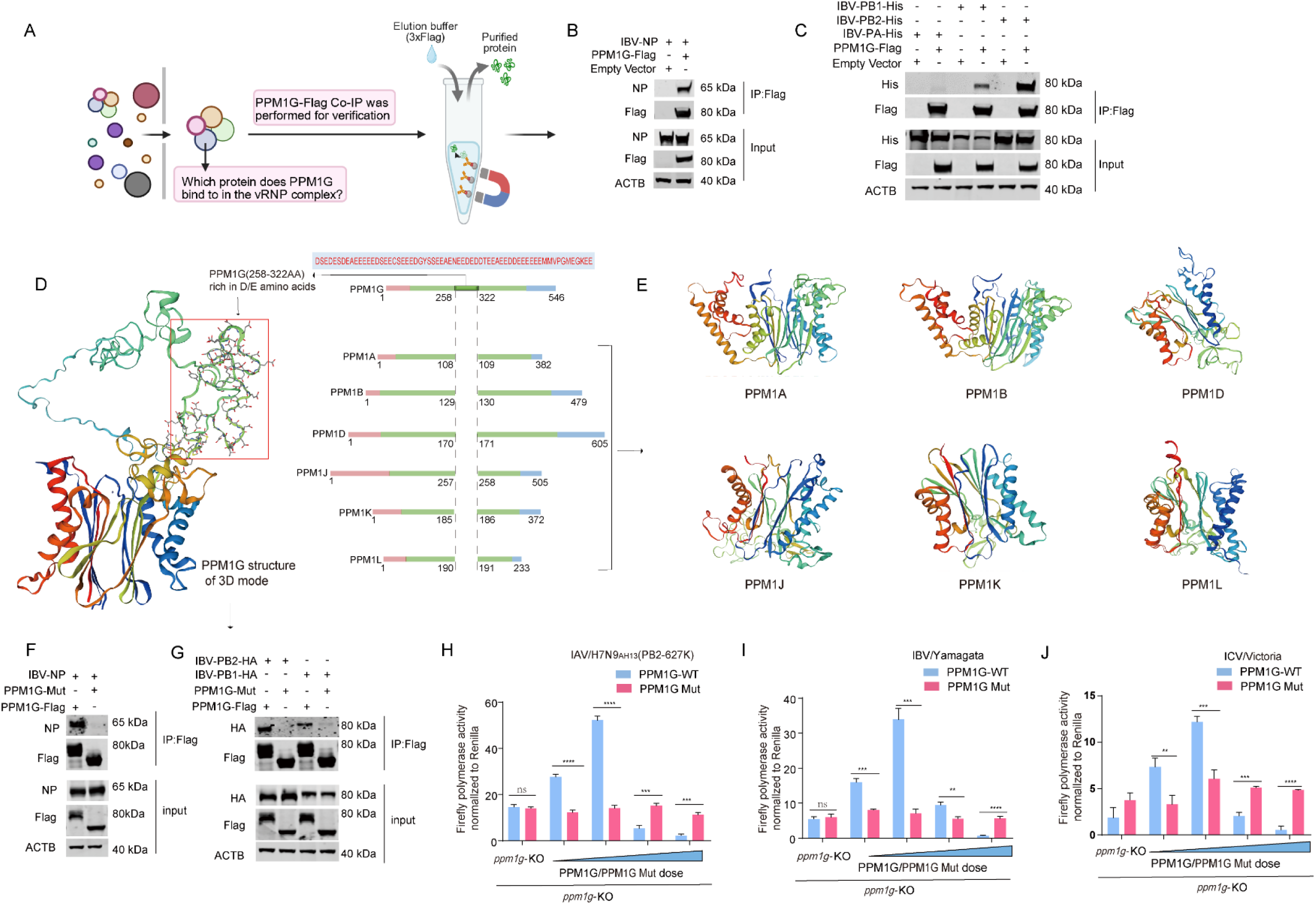
PPM1G binds to viral polymerase and regulates polymerase activity through the PPM1G acidic domain. A-C: Schematic of the Co-IP experiment to investigate potential interactions of PPM1G with vRNP subunits (PA, PB1, PB2, and NP) (A). Each vRNP subunit was individually co-expressed with PPM1G in cells, cell lysates were subjected to Co-IP, and the protein binding was detected by WB (B, C). D: Sequence comparison of PPM1G and other PPM family proteins. E: Comparison of the 3D structure of PPM1G with other PPM family proteins. F, G: Identification of the interactions between PPM1G or PPM1G Mut (PPM1G-Δ258-322) and IBV-NP (F), PB1, PB2 (G) using Co-IP. Representative data from at least two independent experiments are shown. H-J: Minigenome assays of polymerase activity from H7N9_AH13_(PB2-627K) (H), IBV-Yamagata (I), and ICV/Victoria (J) in human HEK293T cells transfected with PPM1G or PPM1G Mut. Error bars represent means ± SD from n = 3 independent biological replicates; Statistical significance was determined by two-tailed unpaired t-test. NS = not significant, ** *p* < 0.01, *** *p* < 0.001.

### 4. PPM1G is a phosphatase of IAV/IBV-NP

The dynamic balance of phosphorylation and dephosphorylation of the NP protein is crucial for maintaining influenza virus replication. Whether PPM1G, as a host phosphatase, plays a role in the dephosphorylation process of NP remains unknown. IAV NP is phosphorylated[17, 35, 36]. However, the phosphorylation of IBV NP protein has been rarely reported. We found that IBV-NP protein formed two bands using Phos-tag™ gel, when stimulated with 12-myristate 13-acetate (PMA 25 µM, PMA can upregulate protein phosphorylation by activating PKC family members and their downstream effectors) and okadaic acid (OA 100 nM, OA is a phosphatase inhibitor)[17], the upper band of NP significantly increased (Figure 6A), cell lysates were treated with the phosphatase phosphatases CIP (calf intestinal phosphatase) and λPP (λ protein phosphatase), after which the upper band of NP was significantly reduced or disappeared completely (Figure 6A). This demonstrated that the band visible above the NP band was its phosphorylated form and confirmed the phosphorylated form of IBV/Yamagata NP. We observed that the presence of PPM1G reduced the overall protein expression levels of vRNP (Figure 6B). IBV-NP was observed to have two bands (Figure 6B lane 1), overexpression of PPM1G significantly decreased the phosphorylation of NP (Figure 6B lane 2, 3), with increasing doses of PPM1G, not only did the phosphorylated NP band diminish, but the protein levels of the non-phosphorylated NP form were also reduced (Figure 6B lane 4). In IBV-infected A549 cells, we also observed that PPM1G reduces the phosphorylation of endogenous NP protein (Figure 6C). These results suggest that PPM1G-mediated dephosphorylation of NP may disrupt vRNP function, potentially contributing to the subsequent reduction in vRNP protein expression. Further experiments showed that overexpression of PPM1G protein significantly inhibited the phosphorylation of NP (Figure 6D), but had no significant effect on PA, PB1, or PB2 (Figure 6E). In addition, we found that the phosphorylated form of IBV/Yamagata NP was significantly increased in *ppm1g*-KO cells, Co-IP analyses using anti p-T/S antibody demonstrated a significant increase in the phosphorylation of NP in *ppm1g*-KO cells, suggesting that the dephosphorylation function of PPM1G targets NP (Figure 6F) and NP is established as a substrate of PPM1G by these results.

**Figure 6:**
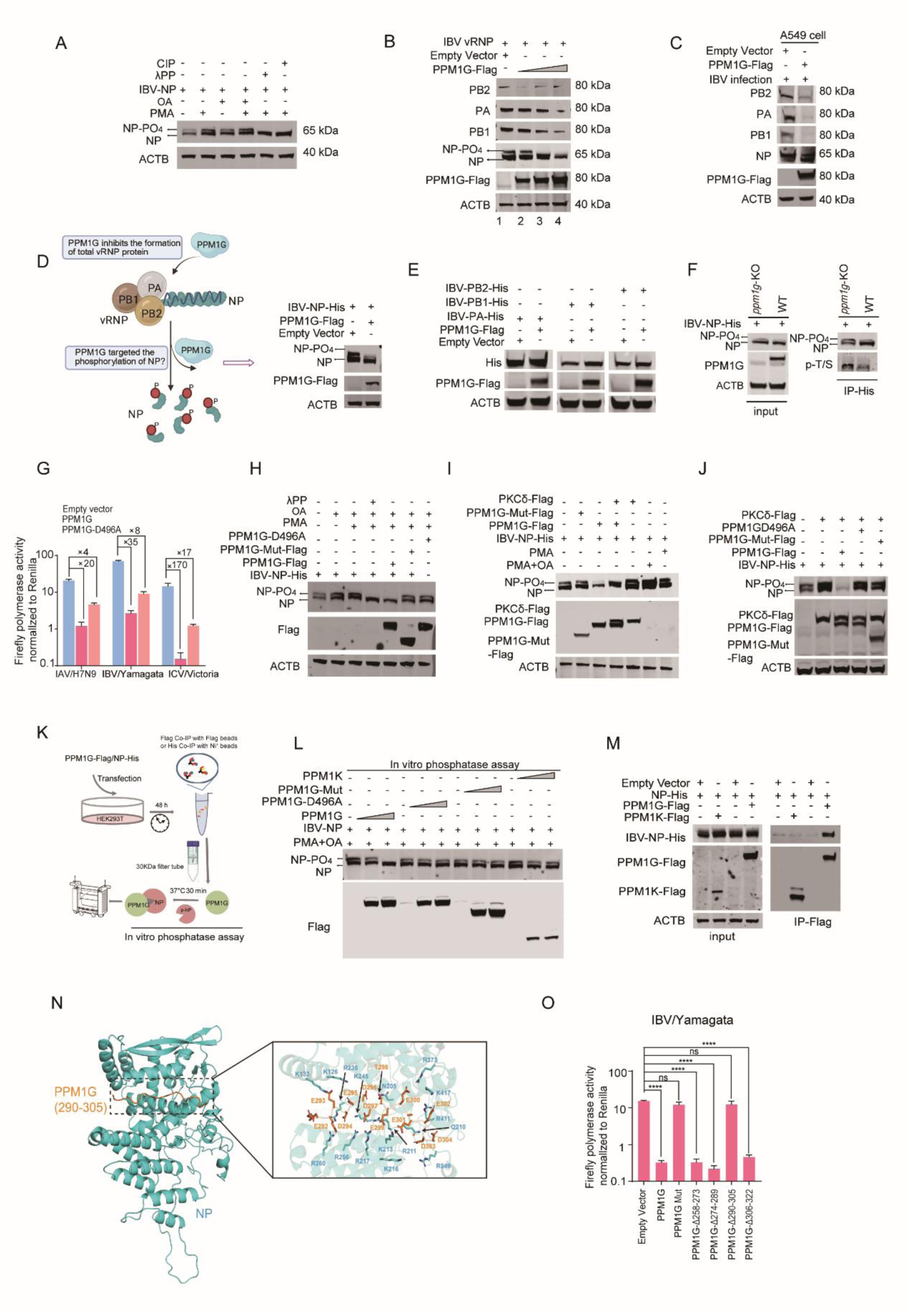
PPM1G is a phosphatase of IBV-NP. A: IBV/Yamagata-NP-His was transfected into HEK293T cells. PMA/OA was added, and the cells were incubated for 2h before cell lysis, after which the cell lysate was either treated for 30 min with CIP or λPP. Polyacrylamide gels with Phos-tag™ gel were used. B, C: A549 cells overexpressing PPM1G-Flag or empty control were transfected with the IBV/Yamagata vRNP (B) or infected with IBV/Yamagata (MOI = 2) (C). After 24 h, the protein content of vRNP in the cell lysate was assessed using WB D-E: PPM1G-Flag was separately co-transfected with IBV/Yamagata NP-His (D), PA-His, PB1-His or PB2-His (E) into HEK293T cells, and changes in protein levels were assessed using WB. F: Detection of phosphorylation of IBV-NP in WT and *ppm1g*-KO cells by Western blotting. G: Minigenome assays of H7N9_AH13_(PB2-627K), IBV-Yamagata and ICV with co-expressed PPM1G-Flag and PPM1G-D496A-Flag. H: NP-His was co-transfected into HEK293T cells together with the empty vector, or with PPM1G- Flag, PPM1G-Mut-Flag or PPM1G-D496A-Flag. The cells were treated with PMA to promote NP phosphorylation. Subsequently, the cell lysate was treated with either CIP or λPP. The change in levels of IBV-NP phosphorylation was assessed using a polyacrylamide gel with Phos-tag. I, J: NP-His was transfected into HEK293T cells (PMA and OA were added) together with PPM1G-Flag, PPM1G-Mut-Flag or PKCδ (I). NP-His and PKCδ were co-transfected into HEK293T cells together with the empty vector, PPM1G-Flag, PPM1G-Mut-Flag or PPM1G-D496A-Flag (J). IBV- NP phosphorylation was assessed using a polyacrylamide gel with a Phos-tag. K-M: IBV-NP-His, PPM1G-Flag, PPM1G-Mut-Flag PPM1G-D496A-Flag and PPM1K-Flag proteins were isolated by Co-IP (PMA and OA were added to cells to promote NP phosphorylation enhancement) and then subjected to a dephosphorylation experiment (see schematic in K)). The phosphorylation modification of NP protein was assessed using WB (L). IBV-NP-His, PPM1G-Flag and PPM1K-Flag were co-transfected into HEK293T cells and then subjected to anti-Flag precipitation (M). All Western blot experiments were independently repeated at least twice with consistent results. N: Predicted molecular docking model of the PPM1G peptide (residues 290-305) in complex with IBV-NP. O: PPM1G Mut, PPM1G-Δ-258-273 mut, PPM1G-Δ-274-289 mut, PPM1G-Δ-290-305 mut or PPM1G-Δ-306-322 mutant proteins were transfected separately into HEK293T cells together with the IBV/Yamagata minigenome reporter system. Luciferase activity was measured 24 h after transfection. Error bars represent means ± SD from n = 3 independent biological replicates; statistical significance was determined by two-tailed unpaired t-test. NS=not significant, **** *p* < 0.0001.

PPM1G-D496 is a key residue reported to be critical for its phosphatase activity[37]. In this study, a PPM1G-D496 mutation (PPM1G-D496A) significantly attenuated the inhibitory effect of PPM1G on the polymerase activities of IAV, IBV and ICV, indicating that the phosphatase activity of PPM1G played a certain role in regulating influenza virus replication (Figure 6G). Subsequently, the results of intracellular experiments further demonstrated that PPM1G was able to dephosphorylate IBV-NP, while PPM1G-mut and PPM1G-D496A could not (Figure 6H). In addition, we found that protein kinase PKCδ can significantly promote the phosphorylation of IBV-Yamagata NP, and PPM1G can dephosphorylate PKCδ-mediated NP phosphorylation (Figure 6I, J). These results give further evidence that PPM1G may be a phosphatase of IBV-NP.

To further verify our hypothesis, an in-vitro phosphorylation assay was performed (Figure 6 K). We found that PPM1G protein could dephosphorylate IBV-NP in vitro (Figure 6L). Consistent with our hypothesis, we also found that PPM1G-D496A does not dephosphorylate IBV-NP in vitro. At the same time, PPM1K was unable to dephosphorylate IBV-NP and could not interact with it (Figure 6L, M), which was consistent with previous research suggesting that PPM1G was the only member of the PPM family able to significantly inhibit influenza virus polymerase activity (Figure 4). These results indicate that PPM1G is a phosphatase of IBV-NP.

The structural prediction of the PPM1G acidic domain and vRNP subunits using ALPHA-fold software revealed that 290-305 could bind to NP protein through salt bridge interactions (Figure 6 N), which was consistent with the results of the polymerase activity experiments (Figure 6O). We found that neither PPM1G-Δ290-305 nor PPM1G-Δ258-322 were able to inhibit the activity of influenza virus polymerases (Figure 6O). We divided the 290-305 domain of PPM1G into three segments or 16 single point mutations, we found that PPM1GΔ290-295, PPM1G-Δ296-300 and mut PPM1GΔ301-305 still inhibit polymerase activity (Figure S6A-C). Single-point mutations of 16 amino acids were performed, and PPM1G still maintained its inhibitory effect on polymerase activity (Figure S6D). These results indicated that the 290-305 domain of PPM1G is the minimum key domain in the regulation of influenza virus replication.

Compared with IBV-NP, the phosphorylated form of IAV-NP was difficult to observe on a polyacrylamide gel and a phosphorylated-tag. Therefore, we used PMA and OA to promote the phosphorylation of IAV-NP to observe the relationship between IAV NP and PPM1G. When PPM1G is overexpressed, it was able to significantly inhibit the overall protein expression of H1N1_WSN_-vRNP (Figure S7A, B). Moreover, PPM1G was able to independently inhibit the phosphorylation of IAV-NP (Figure S7F) without affecting the PA, PB1 or PB2 (Figure S7C-E). This is consistent with the ability of PPM1G to dephosphorylate IBV-NP. PKC δ is a phosphate kinase of IAV-NP and can promote the phosphorylation of IAV-NP. This phosphorylation is inhibited by PPM1G, but not by PPM1G mutants (Figure S7G). These results further demonstrate that PPM1G can dephosphorylate NP from IAV and IBV, possibly serving as a phosphatase to regulate influenza virus replication.

### 5. PPM1G regulates NP protein dephosphorylation by targeting NP-S226/S320/S412/S486

Our findings indicate that PPM1G may serve as a specific phosphatase mediating IBV-NP dephosphorylation. We subsequently identified and characterized the key phosphorylation sites on IBV-NP that are regulated by PPM1G.

IBV-NP protein was purified and subjected to MS analysis at the Institute of Biophysics, Chinese Academy of Sciences to identify the possible phosphorylation sites of NP. We combined 3-5 mass spectrometry analyses to obtain the following 15 possible phosphorylation sites on the NP protein: S50, T55, S58, T73, T127, T163, S167, S188, S320, S327, T406, S412, S459, S465 and S486 (Figure 7A). Here, we extended our investigation to include IBV-NP-S226 and S463[17], two previously reported potential phosphorylation sites that were not detected in our mass spectrometry analysis. We further validated the potential phosphorylation of these sites by using the DTU Healthtech website (https://services.healthtech.dtu.dk/), the locations of these potential phosphorylation sites within NP protein motifs and their predicted scores are presented in Figure 7B. To investigate the role of NP phosphorylation in modulating polymerase function, we proposed substituting phosphorylation sites with alanine (A) to mimic a dephosphorylated state and with aspartic acid (D) to mimic a phosphorylated state. The results showed that the mutation NP-S320A and NP- S226A significantly inhibited the activity of the influenza virus polymerase and the mutation NP- S412A slightly attenuated polymerase activity, indicating that non-phosphorylation of NP at most sites would have compensatory effects on other sites, without affecting the support function of NP to polymerase activity (Figure 7C). The hyperphosphorylation of NP-S226D, NP-S320D, NP- S412D, and NP-S486D significantly impair polymerase activity (Figure 7D). This indicates that the phosphorylation sites NP-S226, NP-S320, NP-S412 and NP-S486 are crucial to the replication of IBV.

**Figure 7:**
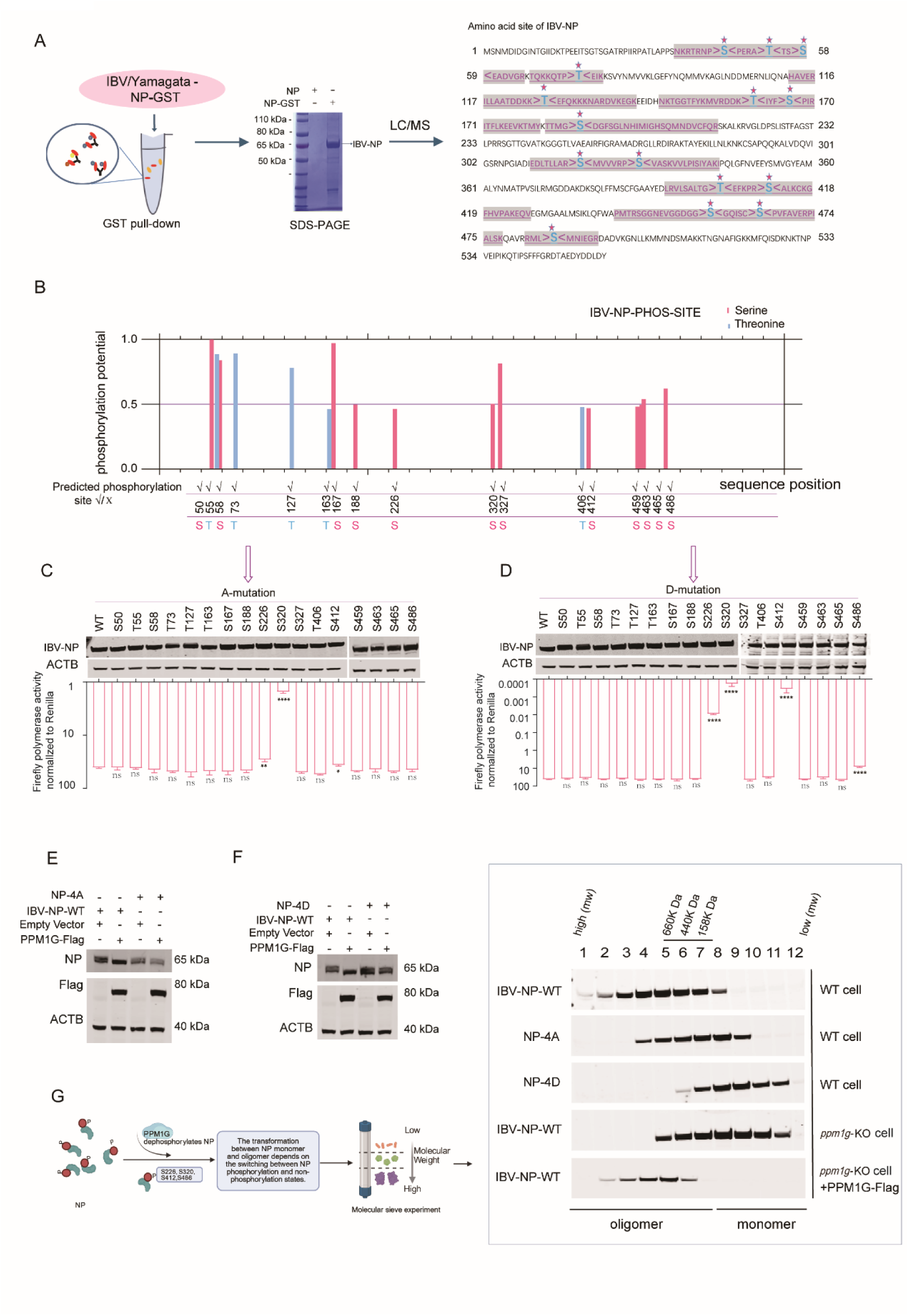
PPM1G targets NP-S226/S320/S412/S486 by regulating the oligomerization of NP. GST-tagged IBV-NP constructs were transfected into HEK293T cells. The pull-down samples resolved by SDS-PAGE and stained with Coomassie blue, NP derived peptides were identified by mass spectrometry (MS) analysis. Fifteen potential phosphorylation sites (S50, T55, S58, T73, T127, T163, S167, S188, S320, S327, T406, S412, S459, S465 and S486) on the IBV-NP protein were identified by mass spectrometry (MS) analysis (A). The DTU Healthtech website (https://services.healthtech.dtu.dk/) predicts that these 17 sites (including S226 and S463 which have been previously reported) may be phosphorylated (B). The scores of the phosphorylation potential of these sites and the positions in NP protein are shown in B. C, D: Minigenome assay of IBV/Yamagata with co-expressed WT-IBV-NP or plasmids encoding IBV-NP with phosphoablative (S/T-to-A) or phosphomimetic (S/T-to-D) mutations. Eerror bars represent the mean ± SD of n = 3 independent biological replicates, NS = not significant, ** *p* < 0.01, **** *p* < 0.0001. E, F: PPM1G-Flag and NP-WT or NP-4A (NP-S226A/S320A/S412A/S486A) (E) or NP-4D (NP-S226D/S320D/S412D/S486D) (F) were transfected into HEK293T cells, and changes in NP phosphorylation were assessed by WB using Phos-tag™ SDS-PAGE. G: IBV-NP-WT, NP-4A or NP-4D were transfected into HEK293T cells, and IBV-NP-WT was transfected into *ppm1g*-KO cells or *ppm1g*-KO cells overexpressing PPM1G. Co-IP was performed using anti-His beads after 36 h. The samples were fractionated by gel filtration chromatography using a Superose 6 Increase 10/300 GL column, and the eluted fractions were subsequently analyzed by Western blotting. Representative results from at least two independent experiments are shown.

Previous experiments demonstrated that PPM1G was able to dephosphorylate NP, and we hypothesized that there might be one or more phosphorylation sites targeted by PPM1G. We found that PPM1G could still dephosphorylate NP-S226A, NP-S320A, NP-S412A or NP-S486A (Figure S8A). We thus speculated that PPM1G dephosphorylated NP at multiple sites. We found that NP-S226/S320A, NP-S226/S320/S486A and wild-type NP could be dephosphorylated by PPM1G (Figure S8B). However, the combined mutants NP-S226/S320/S412/S486A (NP-4A) and NP- S226/S320/S412/S486D (NP-4D) were no longer regulated by PPM1G (Figure 7E, F). The NP-4A mutation greatly reduces NP phosphorylation, which identifies these four amino acids as the key phosphorylation sites (Figure 7E). Our repeated unsuccessful attempts to rescue NP-4A and NP-4D mutant viruses via reverse genetics suggest that these four sites are indispensable for viral replication. The above results suggest that PPM1G targets the phosphorylation sites of NP at S226/S320/S412/S486.

Phosphorylation of the influenza virus NP protein is crucial for maintaining its monomeric state, while the dephosphorylated state may be a prerequisite for NP oligomerization. Our data demonstrates that PPM1G can dephosphorylate both IAV-NP and IBV-NP at specific sites (Figure 6, Figure S7). To investigate whether PPM1G plays a critical role in maintaining NP oligomerization, we employed size-exclusion chromatography and observed that, compared to wild-type NP, both IBV-NP-4A and IBV-NP-4D mutations disrupted the formation of NP oligomeric structures in 293T cells (Figure 7G). These findings indicate that the phosphorylation sites targeted by PPM1G significantly influence the transition between NP oligomerization and monomeric states.

We subsequently observed that NP predominantly existed in a monomeric state in *ppm1g*-KO (Figure 7G), suggesting enhanced phosphorylation of NP in the absence of PPM1G. This finding is consistent with the results shown in Figure 6F. Notably, reconstitution of PPM1G into the knockout cells significantly promoted NP oligomerization (Figure 7G). These results demonstrate that PPM1G facilitates NP oligomerization through dephosphorylation, which may subsequently promote vRNP assembly and ultimately regulate influenza virus replication.

### 6. PPM1G regulates viral replication by degrading NP through the autophagic lysosomal pathway

We observed that PPM1G promotes the oligomerization of NP by dephosphorylating it, and in this process, the overall protein level of NP substantially decreased (Figure 6B, H-J, and Figure 7G). How PPM1G regulates NP protein levels, and whether this is one of the key mechanisms by which PPM1G maintains the functional homeostasis of vRNP complexes, are questions we aim to explore. In Figure 3, we observed significant upregulation of PPM1G protein levels in lung tissues of WSN-infected mice, indicating its strong involvement in influenza virus replication. In Figure 8A, we also detected upregulated PPM1G protein expression in IBV-infected A549 cells, staying step with viral protein expression. We initially discovered that under low-level PPM1G overexpression, NP coexpression with vRNA (specifically NS and M segments) or vRNA exhibited enhanced stability compared to free NP, although dephosphorylation happened (Figure 8B). We detected mRNA of NS and M genes in cell lysates, confirming formation of vRNP complexes (Figure 8C). Furthermore, RNA immunoprecipitation (RIP) assays verified physical packaging of NP with vRNA (Figure 8D), providing additional evidence that vRNA exerts a protective effect on NP protein stability. However, the protective effect of vRNA on NP was attenuated with increasing doses of PPM1G expression (Figure 8E). These results indicate that the reduction in free NP levels in the presence of PPM1G may be associated with its dephosphorylation status.

**Figure 8:**
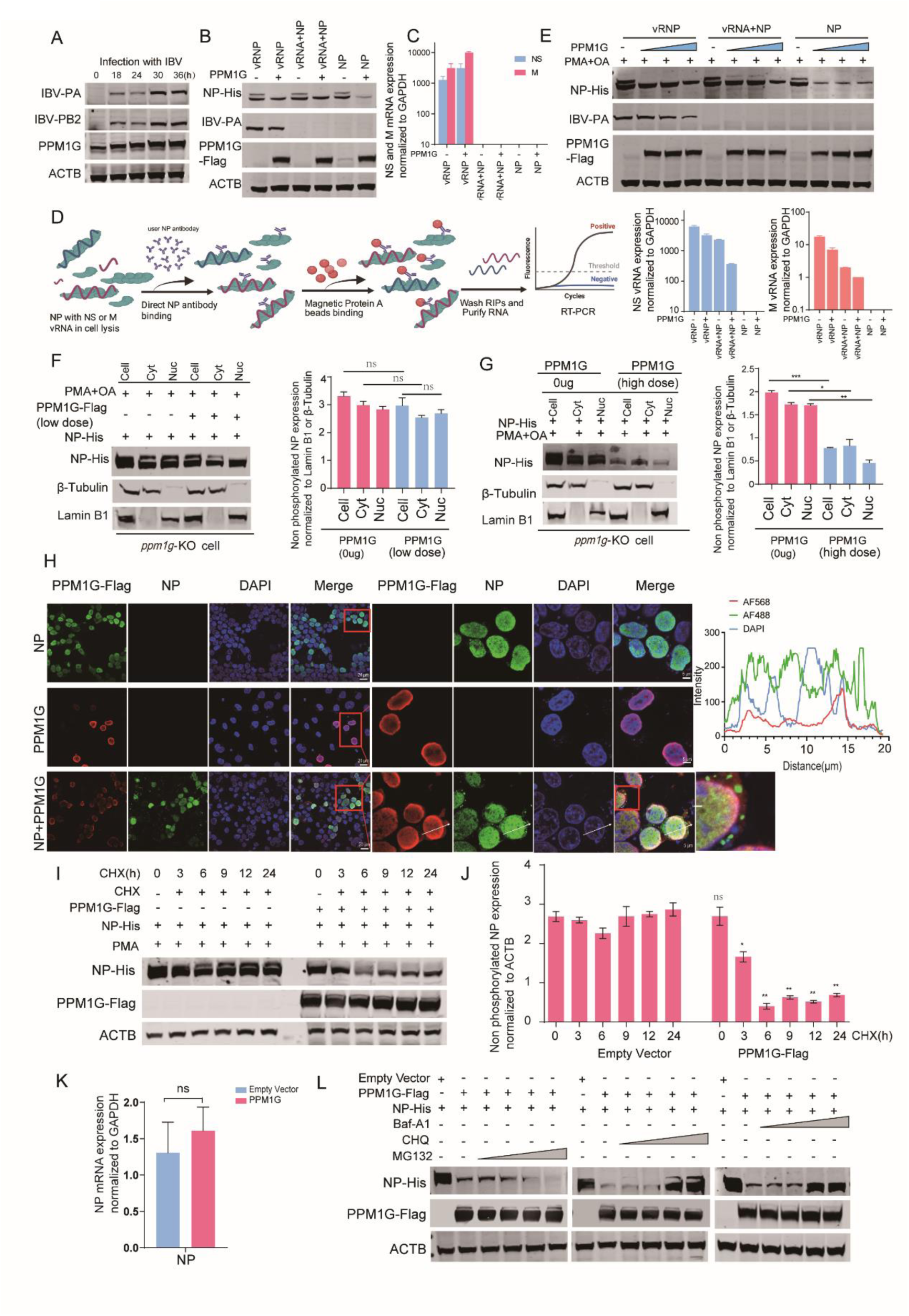
PPM1G overexpression degrades NP through the lysosomal pathway. A-E : A549 cells were infected with IBV (MOI=2), and the expression of IBV viral proteins and PPM1G was detected by WB 36 h later(A). PPM1G was co-expressed with vRNP, vRNA+NP, and NP in cells, respectively. The phosphorylation status of NP was assessed by WB (B); the successful expression of M and NS genes was verified by RT-PCR to confirm the presence of vRNP complexes (C); the interaction between NP and vRNA was validated by RNA immunoprecipitation assay (RIP) (D). Under the same experimental conditions as in B, Western blot analysis was performed to examine both viral protein expression and the phosphorylation status of NP with gradually increasing transfection doses of PPM1G (E). F, G: IBV-NP was co-transfected into *ppm1g*-KO cells with low quantity PPM1G (0.1μg/3.8cm^2^) (F) or high quantity PPM1G (1μg/3.8cm^2^) (G), nuclear and cytoplasmic proteins were extracted using cytoplasmic and nuclear protein extraction kit, and the IBV-NP phosphorylation was assessed using a Phos-tag™ SDS-PAGE. H: Immunofluorescence analysis of the indicated PPM1G-Flag and IBV-NP-His in HEK293T cells, scale bars = 5μm. I-K: IBV-NP-His was co-transfected with empty vector or PPM1G-Flag into HEK293T cells. 24 h after transfection, the cells were treated with CHX (50 μg), and then the cell samples at each time point (0, 3, 6, 9, 12, 24 h) were taken for protein content analysis using WB (I). Quantitative analysis of non-phosphorylated IBV-NP protein from (I) was performed using ImageJ (J). (K) IBV-NP was co-transfected with empty vector or PPM1G-Flag into HEK293T cells. 24 h after transfection, the effect of PPM1G overexpression on the transcription levels of IBV-NP was determined using RT-PCR, error bars represent means ± SD from n = 3 independent biological replicates, statistical significance was determined by two-tailed unpaired t-test, NS = not significant. L: Empty vector or PPM1G-Flag was co-transfected with IBV-NP. 20 h after transfection, 0.5 μL, 1 μL, 1.5 μL, or 2 μL of 10 mM MG132, or 10 mM CHQ or 10 μM Baf-A1 were added separately to 1 mL aliquot of culture medium. Representative data from at least two independent experiments are shown.

We next sought to elucidate how PPM1G modulates NP protein levels. Since vRNP primarily mediates replication functions within the nucleus, while NP exhibits robust nucleocytoplasmic shuttling activity and PPM1G is predominantly localized in the nucleus, we first sought to examine the interaction between PPM1G and NP.

To this end, we performed nuclear-cytoplasmic fractionation assays. In *ppm1g*-KO cells, phosphorylated NP was detected in the nucleus, while phosphorylated forms of NP were significantly reduced at low concentrations of PPM1G (Figure 8F). This further confirmed that PPM1G targets NP in the nucleus. Interestingly, the phosphorylated NP and non-phosphorylated NP in whole cells and cytoplasm were significantly reduced at high concentrations of PPM1G (Figure 8G) and we also found that NP was localized to both the nucleus and cytoplasm. When PPM1G was overexpressed, NP and PPM1G were predominantly localized in the nucleus, but NP aggregation foci were also observed in cytoplasm, which may be associated with protein degradation processes (Figure 8H). Consistent with our assumptions, NP was dephosphorylated when PPM1G was overexpressed, and we subsequently found that non-phosphorylated NP was significantly degraded when cycloheximide (CHX) was added (Figure 8I, J), CHX is an inhibitor of protein biosynthesis. The results also verified that the reduction in NP protein levels was independent of the transcription level (Figure 8 K). The addition of autophagosome-lysosome pathway inhibitors Baf-A1 (Bafilomycin-A1) and CHQ (Chloroquine) was able to restore the inhibitory effect of PPM1G on NP protein. While treatment with MG132, a ubiquitin-proteasome pathway inhibitor, did not (Figure 8L). These results suggest that PPM1G mediated degradation of NP protein in the cytoplasm through the autophagic lysosomal pathway.

### 7. PPM1G regulates viral replication by degrading NP through the ATG7 pathway

Proteins can be targeted to lysosomes through three types of autophagy: microautophagy, macroautophagy and chaperone-mediated autophagy (CMA). CMA requires LAMP2A (Lysosomal Associated Membrane Protein 2A) as a receptor[38–40]. When LAMP2A was silenced by siRNA, PPM1G was still able to reduce the expression of NP (Figure 9A). Macrophage/autophagy is mediated by autophagosomes. PPM1G degradation of NP could be blocked with the addition of 3-MA and LY294002 (3-MA and LY294002 block autophagosome formation by inhibiting PI3K[41]) (Figure 9B). In addition, we observed that PPM1G reduced the expression of P62 (SQSTM1) which is known to act as a specific receptor for autophagy[42] (Figure 9C). LC3 is widely used as a marker of autophagy and is conjugated to the autophagosomal membrane lipid phosphatidylethanolamine (PE) after induction of autophagy[43]. Cytoplasmic type LC3 (LC3-I) will enzymatically hydrolyze a small segment of polypeptide and then convert to membrane type (LC3-II)[44]. PPM1G overexpression resulted in NP degradation and promoted the lipidation of LC3 from L3I to LC3II (Figure 9C). Moreover, the aggregated NP strongly colocalized with LC3-II (a marker of autophagosomes), and the addition of CHQ increased the colocalization of NP and LC3II (Figure 9D). We further demonstrated that PPM1G-promoted degradation of NP protein is dependent on LC3. These results suggest that PPM1G may degrade NP protein through autophagic lysosomal pathways.

**Figure 9:**
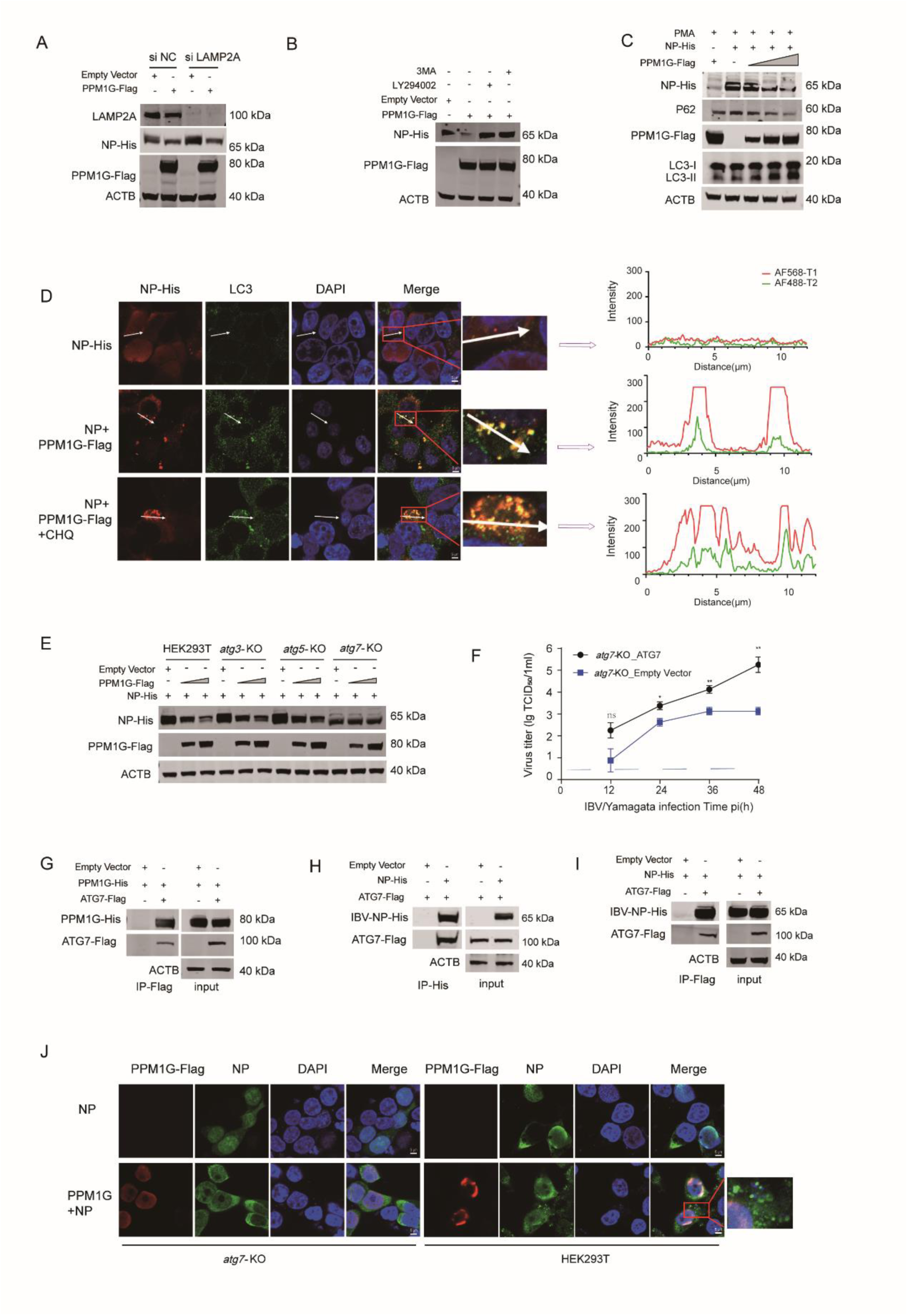
PPM1G degradation of NP is mediated by ATG7 through the autophagy lysosome pathway. A: PPM1G-Flag, *siLAMP2A* and *siNC* were co-transfected with either IBV-NP or empty vector control into HEK293T cells, western blot analysis was performed to examine protein expression. B: PPM1G-Flag was co-transfected with IBV-NP or empty vector into HEK293T cells. At 24 hours post-transfection, the cells were treated with 3-MA (10 mM) and LY294002 (20 μM). C, D: IBV- NP-His was co-transfected with empty vector or PPM1G-Flag into HEK293T cells, and the proteins were subsequently assessed using WB (C). Immunofluorescence analysis of the indicated IBV-NP- His and LC3, scale bars = 5μm (D). E: IBV-NP-His was co-transfected with empty vector or PPM1G-Flag into HEK293T cells, *atg3-* KO cells, *atg5*-KO cells or *atg7*-KO cells, and the proteins were subsequently assessed using WB. F: *Atg7* knockout significantly inhibits IBV/Yamagata influenza virus replication. *Atg7*-KO cells overexpressing ATG7 or empty control were infected with IBV/Yamagata (MOI = 0.01), the virus titers were measured with TCID_50_, error bars represent means ± SD from n = 3 independent biological replicates; Statistical significance was determined by two-tailed unpaired t-test, NS = not significant, \**p*<0.05, ** *p* < 0.01. G: PPM1G-His was co-transfected with empty vector or ATG7-Flag into HEK293T cells. Co-IP was performed with anti-Flag beads; the proteins were assessed using WB. H, I: IBV-NP-His was co-transfected with empty vector or ATG7-Flag into HEK293T cells. Co-IP was performed with anti-Flag beads or anti-His beads, and the proteins were assessed using WB. J: Immunofluorescence analysis of the indicated IBV-NP-His and PPM1G-Flag in *atg7*-KO cells, scale bars=5μm.

Autophagy is a process by which cytoplasmic proteins or organelles are sequestered into vesicles, which then fuse with lysosomes to form autolysosomes and degrade the encapsulated contents. This helps to meet the metabolic needs of the cell itself and allows the renewal of some organelles. There are some important autophagy-related proteins (ATGs) in the autophagosome-lysosome pathway, including Beclin1, ATG3, ATG5, and ATG7 [45]. Our results showed that PPM1G significantly reduced NP protein levels in both *atg3*- and *atg5*-knockout cell lines, but not in *atg7*-knockout cell lines (Figure 9E). These *atg3*-, *atg5*-, and *atg7*-knockout HEK293T cell lines had been previously validated in our studies[46]. Influenza virus replication was significantly inhibited in *atg7* KO cells (Figure 9F). Consistent with our speculations, ATG7 was able to interact with PPM1G (Figure 9G) and IBV-NP (Figure 9H, I). In *atg7* KO cells, overexpression of PPM1G could not cause NP aggregation in the cytoplasm (Figure 9J). All these findings demonstrated that overexpression of PPM1G degrades NP protein through the ATG pathway in the cytoplasm, thereby regulating virus replication.

## Discussion

Acting as an “armored train”, the influenza NP protein encapsidates the viral RNA genome. It dynamically orchestrates the genome’s nuclear trafficking to ensure replication within the correct compartment, a process entirely dependent on NP-NP self-assembly. The conversion between the oligomeric and monomeric states of the influenza virus NP protein is a prerequisite for vRNP to perform its replication and transcription, the NP protein monomeric state is regulated by the host’s phosphorylation kinase, and the specific mechanism of NP protein oligomerization regulation is a long-standing unsolved scientific problem. Although Protein phosphatase 6A, (PP6A)[47] and cell division cycle 25 B (CDC25B) phosphatase are probably associated with the dephosphorylation of vRNP or NP[48], the discovery and confirmation of the presence of an NP phosphatase in IAV and IBV have not been discussed accurately or in detail in the past. By comparing analysis of mass spectrometry data between purified viral polymerase and vRNP complexes, we identified PPM1G as the phosphatase for NP protein, while also verifying PPM1G’s dephosphorylation function using IAV and IBV systems.

We first discovered that PPM1G plays a crucial role in regulating influenza virus replication. Both overexpression and knockout of PPM1G significantly reduce the polymerase activity of influenza A, B, and C viruses *in vitro*. Lung-specific conditional PPM1G knockout mice exhibited significantly improved survival rates upon influenza A and B virus infection, demonstrating that appropriate expression levels of PPM1G protein are essential for maintaining the efficient operation of the vRNP replication machinery and viral replication. We further identified that no other members of the PPM family, PPM1G is the phosphatase capable of dephosphorylating NP of both influenza A and B viruses, thereby regulating viral replication. In this study, PPM1G, functioning as a phosphatase for NP, was able to counteract the effect of PKC-mediated upregulation of NP phosphorylation, suggesting that both play a collaborative role in maintaining the dynamic balance of NP phosphorylation.

Different phosphorylation states determine whether NP polymerizes. Dephosphorylation enables NP to undergo oligomerization. The structure of IAV and IBV NP comprise three parts, a head, body and a flexible tail-loop[13, 49].During oligomerization, a change in the conformation of the NP tail loop may promote the interaction of the loop with another NP binding groove, resulting in oligomerization. Such oligomerization is indispensable for both vRNP structural integrity and viral replication. Disassociated NP is typically phosphorylated. NP continuously dissociates from the template strand during transcription and replication and is loaded onto nascent vRNA/cRNA during replication, this process requires NP to switch between monomers and oligomers to assemble vRNP[10, 16, 17, 50]. As expected, we observed that PPM1G promotes the oligomerization of NP protein. Our results illustrate that PPM1G can dephosphorylate NP to promote NP polymerization and then may promote viral replication.

Influenza viruses hijack the host protein PPM1G to execute their biological functions. Our observations demonstrate significantly upregulated PPM1G protein expression in lung tissues of influenza-infected mice and A549 cells, confirming the viral requirement for PPM1G as its key phosphatase to promote NP oligomerization and subsequent viral replication. The upregulation of PPM1G expression represents a cellular response to rapid and extensive viral replication. This leads to the dephosphorylation of a large amount of NP protein. However, such an excess of dephosphorylated NP may be detrimental to viral replication. Through its interaction with ATG7, PPM1G likely triggers a degradation switch, directing NP for elimination via the autolysosomal pathway. This suggests that PPM1G plays a critical role in inducing NP degradation, which may represent an alternative fate for surplus NP within the cell. Notably, similar antiviral mechanisms exist for other pathogens, where autophagosomes can encapsulate viruses to inhibit infection[52–54]. Nevertheless, the regulatory mechanisms underlying PPM1G upregulation require further investigation.

The PPM family contains a series of metal-dependent phosphatases that act as a single subunit enzyme and bind Mn^2+^ or Mg^2+^ [26]. In this study, we identified that PPM1G contains a unique acidic repeat sequence (258-322, rich in Asp/Glu residues) distinct from its family members, which serves as a critical motif for both regulating influenza virus activity and mediating NP binding, targeting this specific sequence with competitive small-molecule peptides or inhibitors could represent a promising therapeutic strategy for anti-influenza drug development. Other studies have demonstrated that PPM1G, through its acidic amino acid domain, inhibits the assembly of P-TEFb into the 7SK snRNP complex, thereby promoting transcriptional elongation in HIV (Human Immunodeficiency Virus), this mechanism distinctly differs from that of PPM1A[55]. Our results demonstrate that PPM1G is involved in the regulation of influenza virus replication through binding of its acidic domain to the substrate, in a mechansim based on its phosphatase activity, this is consistent with the traits of the single subunit PPM1G enzyme. PPM1G’s phosphatase activity is exploited by Kaposi’s sarcoma-associated herpesvirus (KSHV) to function as a negative regulator of innate immune signaling pathways[56].

The phosphorylation of IAV NP plays an important role in regulating virus replication. The NP functions as a phosphorylated protein, with multiple sites on IAV-NP undergoing phosphorylation, the phosphorylation of S407 and S165 blocks the oligomerization of NP, inhibiting the assembly of the vRNP complex and therefore also genome replication[17–19]. Mutants of IAV NP (E339A, R416A and S486A) can also affect the form of NP oligomerization[57, 58]. Phosphorylation of sites S226 and S463 of IBV NP blocks virus growth by inhibiting the formation of NP oligomers [17, 59]. In this study, the S226/S320/S412/S486 phosphorylation sites in IBV-NP protein serve dual critical roles: as primary targets for PPM1G-mediated dephosphorylation and as key regulators of influenza viral polymerase activity. These findings establish a structural foundation for developing targeted anti-influenza therapeutics.

Based on our data, we propose a model elucidating the role of PPM1G in regulating influenza virus replication. The phosphorylation process is dynamic and reversible, PPM1G was involved in the conversion between the phosphorylated and dephosphorylated states of IBV-NP in a process related to PKCδ (Figure 10-(1)). PPM1G dephosphorylates NP to promote its oligomerization, thereby facilitating vRNP assembly and genome replication (Figure10-(2)). Dephosphorylated NP undergoes PPM1G-mediated degradation via the ATG7-dependent autolysosomal pathway (Figure 10-(3)).

**Figure 10:**
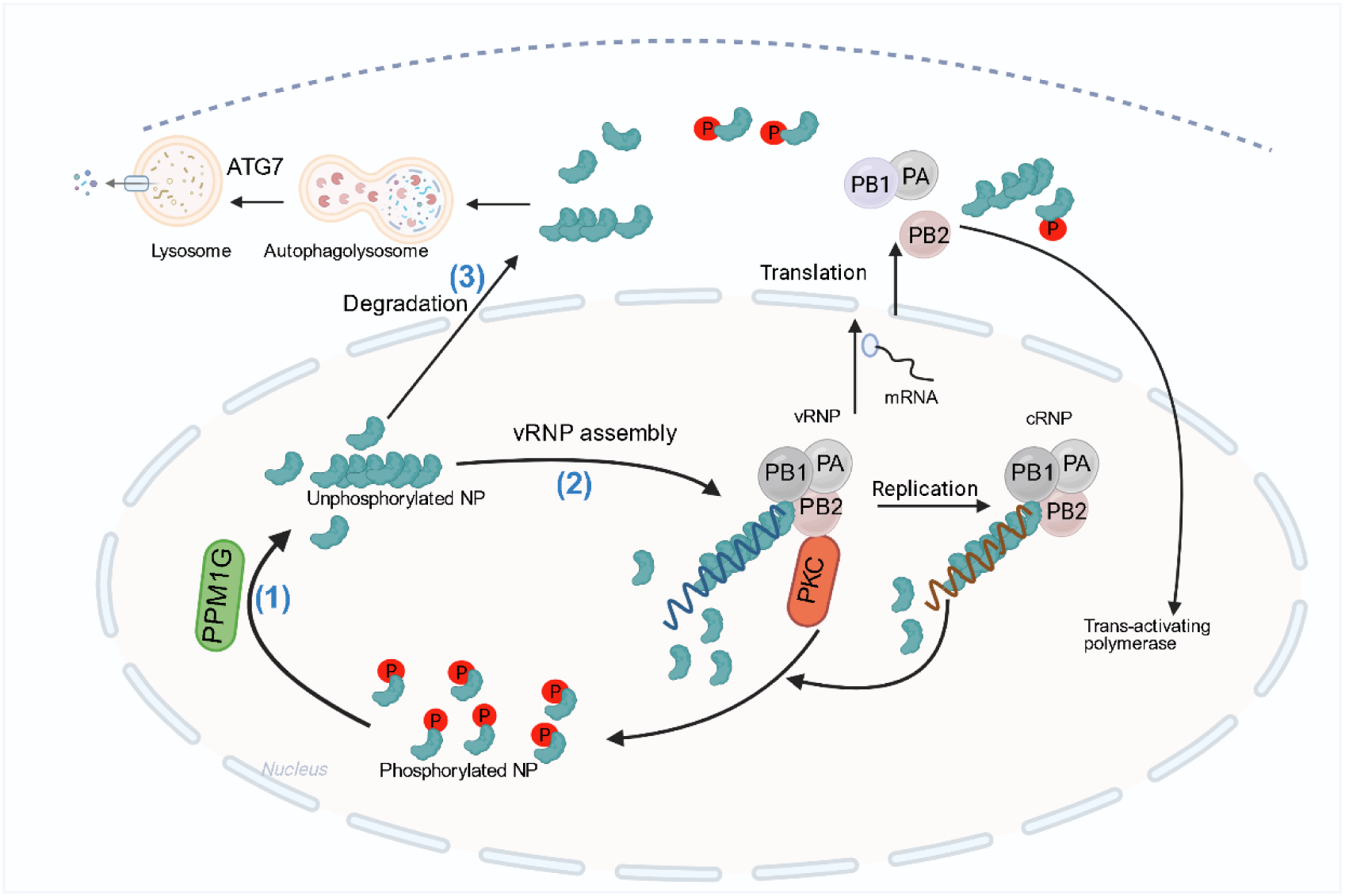
Schematic illustration showing the mode of PPM1G regulation of influenza virus replication. (1) The influenza virus NP protein can exist in a monomeric or oligomeric state. A reversible phosphorylation process regulates the transition of NP monomers to oligomers. PPM1G, a host phosphatase, can dephosphorylate the viral NP protein, thereby promoting the oligomerization of NP protein. The host protein kinase PKC can phosphorylate NP and promote the production of NP monomers. (2) The NP protein dephosphorylated by PPM1G is likely to form oligomeric forms, which may facilitate the assembly of vRNP and further promote replication of the influenza virus genome. (3) NP protein aggregation is promoted by PPM1G, and NP protein is then degraded through the ATG7-mediated autophagy lysosome pathway, ultimately leading to a decrease in influenza virus replication. This figure was created with BioRender (https://www.biorender.com).

In our study, we explain how PPM1G regulates the efficient functioning of vRNPs by modulating its own expression levels, thereby maintaining vRNP homeostasis. The dual role of the host protein PPM1G in regulating influenza virus replication was defined and revealed in detail in this study. These findings offer deeper insights into how host factors influence influenza virus replication, thereby aiding the development of novel anti-influenza agents.

## Materials and Methods

### Cells, Viruses, and Plasmids

Human embryonic kidney 293T cells (HEK293T), Madin-Darby canine kidney cells (MDCK), human lung adenocarcinoma epithelial cells (A549), *ppm1g*-knockout HEK293T cell lines (*ppm1g-* KO), *ppm1g* stable knockdown cells and stably expressing Flag-tagged PPM1G cells (*ppm1g*+/+) were maintained in Dulbecco’s Modified Eagle Medium (DMEM; Sigma) supplemented with 10% fetal bovine serum and 1% penicillin and streptomycin. The IAV/H1N1_WSN_ virus (titer of 10^5.5^ TCID_50_/0.1 mL) and IBV/Yamagata virus (titer of 10^5^ TCID_50_/0.1 mL) used in this study were maintained in the laboratory. Plasmids encoding the polymerases of IAV and IBV were generated as previously described and were taken from lab stocks. The other pCAGGS expression plasmids encoding *SET* (NM_001122821.2, NP_001116293.1), *STOML2* (NM_001287031.2, NP_001273960.1), *SSRP1* (NM_003146.3, NP_003137.1), *PPM1G* (NM_177983.3, NP_817092.1), mutant plasmids of pCAGGS-PPM1G-Δ258-322-Flag, pCAGGS-PPM1G-Δ258-274-Flag, pCAGGS-PPM1G-Δ275-289-Flag, pCAGGS-PPM1G-Δ290-305-Flag, pCAGGS- PPM1G-Δ306-322-Flag, pCAGGS-PPM1G-Δ290-295-Flag, pCAGGS-PPM1G-Δ296-300-Flag, pCAGGS-PPM1G-Δ301-305-Flag and pCAGGS-PPM1G-D496A-Flag were constructed for this project. *PPM1A* (NM_021003.5, NP_066283.1), *PPM1B* (NM_001033557.3, NP_001028729.1), *PPM1D* (NM_003620.4, NP_003611.1), *PPM1J* (NM_005167.7, NP_005158.5), *PPM1K* (XM_017007804.2, XP_016863293.1), *PPM1L* (NM_001317911.2, NP_001304840.1), *PKCδ* (NM_001310682.1, NP_001297611.1) and *ATG7* (NM_001349232.2, NP_001336161.1) were pur-chased from Miaoling Biotechnology Co., Ltd (Wuhan, China).

The IBV/Yamagata NP mutations of pCAGGS-IBV-NP S50A/D, pCAGGS-IBV-NP T55A/D, pCAGGS-IBV-NP S58A/D, pCAGGS-IBV-NP T73A/D, pCAGGS-IBV-NP T127A/D, pCAGGS- IBV-NP T163A/D, IBV-NP T167A/D, pCAGGS-IBV-NP S188A/D, pCAGGS-IBV-NP S226A/D, pCAGGS-IBV-NP S320A/D, pCAGGS-IBV-NP S327A/D, pCAGGS-IBV-NP T406A/D, pCAGGS-IBV-NP S412A/D, pCAGGS-IBV-NP S459A/D, pCAGGS-IBV-NP S463A/D, pCAGGS-IBV-NP S465A/D, pCAGGS-IBV-NP S486A/D, pCAGGS-IBV-NP S226, S320,S412,S486A/D and the 16 single point mutants of the 290-305 region of PPM1G were constructed by single point mutation.

### *Ppm1g* Knockout Cell Lines

The gRNAs design tool (http://crispor.tefor.net/) was used to design the sgRNA1 (5’- CAACACGGTGAAGTGCTCCG-3’), and the sgRNA2 (5’-GCCATGTTTTCTGTCTACGA-3’) of *ppm1g*. The designed sgRNA sequence was inserted into the pMD-18T vector. SeqBuilder software was used to design the primers. The sgRNA1 primers were: 5’-TGTGGAAAGGACGAAACAC- CGCATGAGAAGACTATCGAAC-3’, 5’-ATTTCTAGCTCTAAAACCGGAGCACTTCAC-CGTGTTGC-3’, and the sgRNA2 primers were: 5’-TGGAAAGGACGAAACAC- CGGCCATGTTTTCTGTCTACGA-3 ’, 5’-CTATTTCTAGCTCTAAAACTCGTAGA-CAGAAACATGGC-3’. Plasmid extraction was performed after successful sequencing. Briefly, HEK293T cells were co-transfected with 1µg pMJ920 plasmid (obtained from Jennifer Doudna) and 1µg sgRNAs into one well of a 6-well plate by PEI. 24 h after transfection, GFP-positive cells were sorted by fluorescence-activated cell sorting (FACS), and monoclonal knockout cell lines were then screened using WB.

### *Ppm1g* Stable Knockdown Cell Lines

In this study, the pLVshRNA-EGFP (2A) puro vector was used to construct the *ppm1g* shRNA interference stable cell line. First, the sh-*ppm1g* stable knockdown lentiviral expression vector was constructed. 0.9 µg pcGP, 0.2 µg Pvsv-G, and 0.9 µg pLV shRNA-EGFP (2A) puro-*ppm1g* -sh1 or pLV shRNA-EGFP (2A) puro-*ppm1g*-sh2 were co-transfected into cells using PEI. After 48 h, the cell supernatant was collected, and the cell debris was removed by centrifugation, leaving the pseudovirus supernatant. HEK293T cells were then infected with the collected pseudovirus supernatant, and flow sorting was performed. The cells with green fluorescence were sorted into 96 well plates, and the cells were cultured and were subsequently identified using WB.

### *Ppm1g* Stable Overexpression Cell Lines

The recombinant plasmid pLPCX-PPM1G-Flag was constructed using homologous recombination. 0.9 µg pcGP, 0.2 µg Pvsv-G and 0.9 µg pLPCX-PPM1G-Flag were co-transfected into HEK293T cells using PEI. After 48 h, the cell supernatant was collected, and the cell debris was removed by centrifugation, leaving the pseudovirus supernatant containing pLPCX-PPM1G-Flag. When the density of HEK293T cells reached 50%, 1 mL of pLPCX-PPM1G-Flag pseudovirus supernatant was mixed with 1 mL of cell culture medium and added to a 6-well plate. Groups of transfected cells were then grown in medium containing different concentrations of puromycin (0, 0.25, 0.5, 0.75, 1, 1.5, and 2 µg/mL). The culture medium was changed after 48 h, and the growth status of the cells was recorded every day. The culture medium was replaced every 2 days. After the cells in the negative control group were all dead, the surviving puromycin resistant cells were sorted into 96 well plates using flow cytometry, and the expression of the PPM1G-Flag protein was assessed.

### Mass Spectrometry

HEK293T cells were transfected with three experimental groups: Group A (Flag-tagged RdRp), Group B (Flag-tagged vRNP), and Group C (untagged vRNP as a control). After Co-IP with Flag magnetic beads, samples were separated using SDS-PAGE and were stained with Coomassie brilliant blue. Other steps referred to previous studies[32].

### Mice and mice infection

Lung-specific *ppm1g*-deficient (*ppm1g* ^flox/flox^-Sftpc cre) mice and *ppm1g* ^flox/flox^ mice were purchased from Cyagen (Suzhou, China) and bred in Biosafety Level II laboratory (BSL- II). Wild type (WT) C57BL/6J mice were purchased from Changsheng Biotechnology Co., Ltd. (Liaoning, China). Mice were housed under specific pathogen-free conditions and were used for experiments at 5 to 6 weeks of age.

The mice were anesthetized and inoculated with 10 MLD_50_ of IAV/H1N1_WSN_ or IBV/Yamagata-MD virus through the nasal cavity. Body weight and survival rate were monitored daily. The lung tissue homogenate was prepared by homogenizing fresh frozen lung tissues in PBS for twice (20 seconds each) and then centrifuging at 2500 g for 20 min. TCID_50_ assays were performed using MDCK cells and TCID_50_ was calculated as previously described[32].

### Immunoprecipitation and Western blotting (WB)

Lysis buffer (50 mM Hepes-NaOH [pH 7.9], 100 mM NaCl, 50 mM KCl, 0.25% NP-40, and 1 mM DTT) was used to lyse the cells. The supernatant was incubated with 15 μL of anti-Flag M2 Magnetic Beads (SIGMA-ALDRICH, M8823), or 30 μL of anti-His Magnetic Beads (MCE, HY- K0209) at 4°C for overnight and washed 3-5 times with phosphate-buffered saline (PBS) (for anti-Flag) or 40mM imidazole solution (for anti-His).Then, 100 μL of 3×Flag peptide or 500mM imidazole solution was added and the whole was eluted at 4°C for 20 min or was boiled in 1×loading buffer at 98°C for 8 min. WB was carried out using the following primary antibodies: PPM1G rabbit polyclonal (A20959, Abclonal), Flag mouse monoclonal (F1804, Sigma-Aldrich), Flag rabbit polyclonal (F7425, Sigma-Aldrich), ACTB rabbit monoclonal (AC026, Abclonal), His mouse monoclonal (66005-1, Proteintech), ATG7 rabbit polyclonal (A0691, Abclonal), LC3B rabbit monoclonal (A19665, Abclonal), LAMP2A Rabbit monoclonal (ab125068, abcam), rabbit anti-influenza B virus NP (GeneTex, GTX128538) and anti-IAV-NP (GeneTex, GTX636318). IAV-PB1, IAV-PB2, IAV-PA, IBV-PB1, IBV-PB2 and IBV-PA are monoclonal antibodies generated in our lab.

### Cell viability assay

HEK293T cells were seeded in 96 well plates at a density of 1000 cells well/200 μL medium 24 h before the experiment. After treatment, cell viability was analyzed using the CCK8 reagent (Celling Kit-8).

### Immunofluorescence assay

The cells were transfected or treated with drugs according to the experimental conditions, and were then fixed with 4% paraformaldehyde (Beyotime, China) at room temperature for 15 min. They were then washed 3 times with 1× PBS containing 1% Triton X-100 and incubated for 10 min. The cells were then incubated with 3% skim milk at room temperature for 2 h. The cells were subsequently incubated overnight at 4 °C with primary antibodies (rabbit anti Flag antibody 1:300; rabbit anti-NP 1:200; rabbit anti-LC3 1:200; mouse anti-His 1:200), then washed 3 times with 1× PBS. Cells were incubated with Alexa Fluor-conjugated secondary antibodies (488 or 568 goat antimouse/rabbit IgG; Invitrogen) for 1 h at room temperature. Nuclei were counterstained with 4’,6- diamidino-2-phenylindole (DAPI) for 10 min, and images were acquired using a Zeiss LSM 880 confocal microscope.

### Influenza virus infection

IAV and IBV were rescued from a 12-plasmid rescue system. HEK293T cells were transfected with 1 µg each of eight pPolI plasmids and 2 µg each of pCAGGS-NP, pCAGGS-PA, pCAGGS- PB1, and pCAGGS-PB2. The virus was propagated in embryonated eggs for 48 h to form the virus stocks used to infect the cells. The cells were washed twice with 1×PBS and incubated with the virus for 2 h. Cells were then cultured at 37°C in DMEM containing 0.3% BSA and TPCK (N-Tosyl-L-phenylalanyl chloromethyl ketone)-trypsin at 0.2-1 µg/ml. At the indicated time points, the culture supernatant was harvested, and the viral titers in the cells were determined using standard methods. The NP content in the supernatants was measured using an antigen capture enzyme-linked immunosorbent assay (Ac-ELISA) from our lab.

### RNA Extraction, Reverse Transcription and Fluorescence Quantitative PCR

Total RNA was extracted from the HEK293T cells using RNeasy mini kit (Qiagen). The reverse transcription kit (Cat. RR047A, Takara, China) was used for reverse transcriptase. The 20 µL reverse transcription reaction system was established using random primers and reverse transcriptase to synthesize cDNA. 1 µg of RNA from each sample was used cDNA synthesis. SYBR Premix Ex Taq II (Tli RNaseH Plus) kit was used for fluorescent quantitative reverse transcription PCR (qRT-PCR). The reaction system was as follows: forward/reverse primer (10 mM) 1 µL, TB Green Premium Ex Taq II 10 µL, ddH_2_O 6 µL, cDNA 2 µL. The mixed samples were detected using Agilent Mx 3005P fluorescence quantitative PCR. Signals were collected according to the manufacturer’s instructions following the two-step PCR amplification reaction. After the reaction, the specificity of the amplified product was analyzed through the dissolution curve. The PCR procedure was as follows: 95 °C for 10 min; 95 °C for 30s, 60 °C for 60s, and then 40 cycles of 95 °C 60s, 55 °C 30s, 95 °C 30s. *Ppm1g* gene specific primers (5’-GCTGCATGAAGAGGCTACCA-3’ and 5’-TGTCCCACCTCCAGATTTGC-3’), *IBV-NP* gene specific primers (5’-ATGAAGTAGGTG-GAGACGGAG-3’and 5’-AGCCATTGAATCATTCATCATC-3’), *IAV-NP* gene specific primers mRNA (5’-CGATCGTGCCCTCCTTTG-3’;and 5’-CCAGATCGTTCGAGTCGT-3’). *GAPDH* gene specific primers (5’-GAAGGTGAAGGTCGGAGTC-3’and 5’-GAAGATGGTGATGGGATTTC-3’) were used. The 2^−ΔΔ^Ct method was used to determine the relative mRNA expression of genes normalized by GAPDH.

### Dosing treatment experiment

20 h following transfection of HEK293T cells, 1 mL aliquots of cell culture were taken and 0.5 μL, 1 μL, 1.5 μL, or 2 μL of 10 mM MG132, 10 mM CHQ, or 10 μM BAF-A1 were added to each 1 mL to form three different concentration gradients. Samples were supplemented with DMSO to achieve the correct concentration. After 8-10 h of treatment with the drug, 200 μL cell lysates were prepared (containing 1× protease inhibitor and phosphatase inhibitor), were lysed at 4 °C for 10 min, and were then centrifuged at 10,000 g at 4 °C for 5 min. Samples were prepared by mixing 100 µL supernatant with 25 µL 5× SDS loading buffer. The samples were boiled at 100 °C for 8 min before WB.

Phosphorylation of NP was stimulated with 2.5 μM PMA or 100 nM OA. After treatment, the cells were placed in the incubator for 2 h, and samples were then collected.

24 h after transfection of HEK293T cells, 100 μL of cell lysate containing 1× protease inhibitors and phosphatase inhibitors were added to each well of a 12-well plate. After lysis, the cells were centrifuged at 10,000 g for 5 min at 4°C, and supernatant was added into λPP (λ protein phosphatase) or CIP at 30 °C or 37 °C for 30 min. After the reaction was completed, the reactions were boiled at 100 °C for 10 min and WB was performed immediately afterwards.

### In vitro dephosphorylation assay

HEK293T cells were transfected with PPM1G-Flag, PPM1G-mut-Flag, PPM1G-D496A-Flag, PPM1K-Flag and IBV-NP-His. Flag and his magnetic beads were used separately for Co-IP, and corresponding volumes of PPM1G-Flag, PPM1G-D496A-Flag, PPM1K-Flag and IBV-NP-His protein solutions were obtained. 4 mL of 1× PBS was added to each sample to dilute the protein, and a centrifugal filter (30 kDa) was used to concentrate the protein and replace buffer. After centrifugation at 7,000 g for 30 min at 4 °C, samples were collected for concentration determination and were stored in a refrigerator at -80 °C.

An in vitro dephosphorylation assay was subsequently performed. Purified PPM1G-Flag, PPM1G-mut-Flag, PPM1G-D496A-Flag, PPM1K-Flag and IBV-NP-His were added to phosphatase buffer (1 mM EGTA, 25 mM MgCl_2_, 2 mM DTT, 0.1 % BSA and 250 mM imidazole) and incubated at 30 °C for 40 min. After the reaction, the samples were mixed with 5× SDS loading buffer and boiled before WB analysis.

### Polymerase activity assay

In this study, influenza virus RdRp (PB1, PB2 and PA), NP, vLuc (firefly luciferase gene) and pRL TK (sea kidney luciferase) gene expression plasmids were co-transfected into the minigene reporter system to analyze the polymerase activity. To determine the effect of target protein on influenza virus polymerase activity, PB1 (20 ng), PB2 (20 ng), PA (10 ng), NP (40 ng), vLuc (40 ng) and 2.5 ng TK expression plasmids were co-transfected into a well of HEK293T in a 24 well plate. The double luciferase kit (Promega) was used to investigate the interaction activity between the enzyme and substrate. Firefly luciferase activity was measured using a CentroXSL B960 photometer (Berthold Technologies). All experiments were conducted independently at least twice.

### Gel filtration chromatography

HEK293T or *ppm1g*-KO cells were transfected with IBV-NP-His or IBV-NP-4D-His or IBV- NP-4A-His. The cells were purified using Co-IP with His beads. After eluting the His beads with 500mM imidazole, replacement buffer was carried out with 1×PBS and centrifuged at 13,000 g for 10min 3 times. About 600 μL of the resulting protein was then loaded onto the Superose 6 Increase 10/300 GL (GE healthcare) that pre-equilibrated with 1×PBS. The gel filtration calibration kit HMW (28403842, GE healthcare) was used as molecular weight markers the protein.

### Quantification and statistical analysis

Image J software was used to measure the band intensity from the WB. GraphPad Prism was used for statistical tests. Statistical differences were assessed using an unpaired two-tailed Student’s t-test followed by a Dunnett’s test. Statistical parameters are reported in the figures and figure legends. Error bars represent SEM (standard error of the mean). NS, not significant (*p* > 0.05), \**p* < 0.05, \*\**p* < 0.01, \*\*\**p* < 0.001, \*\*\*\**p* < 0.0001.

---

**Figure S1:**
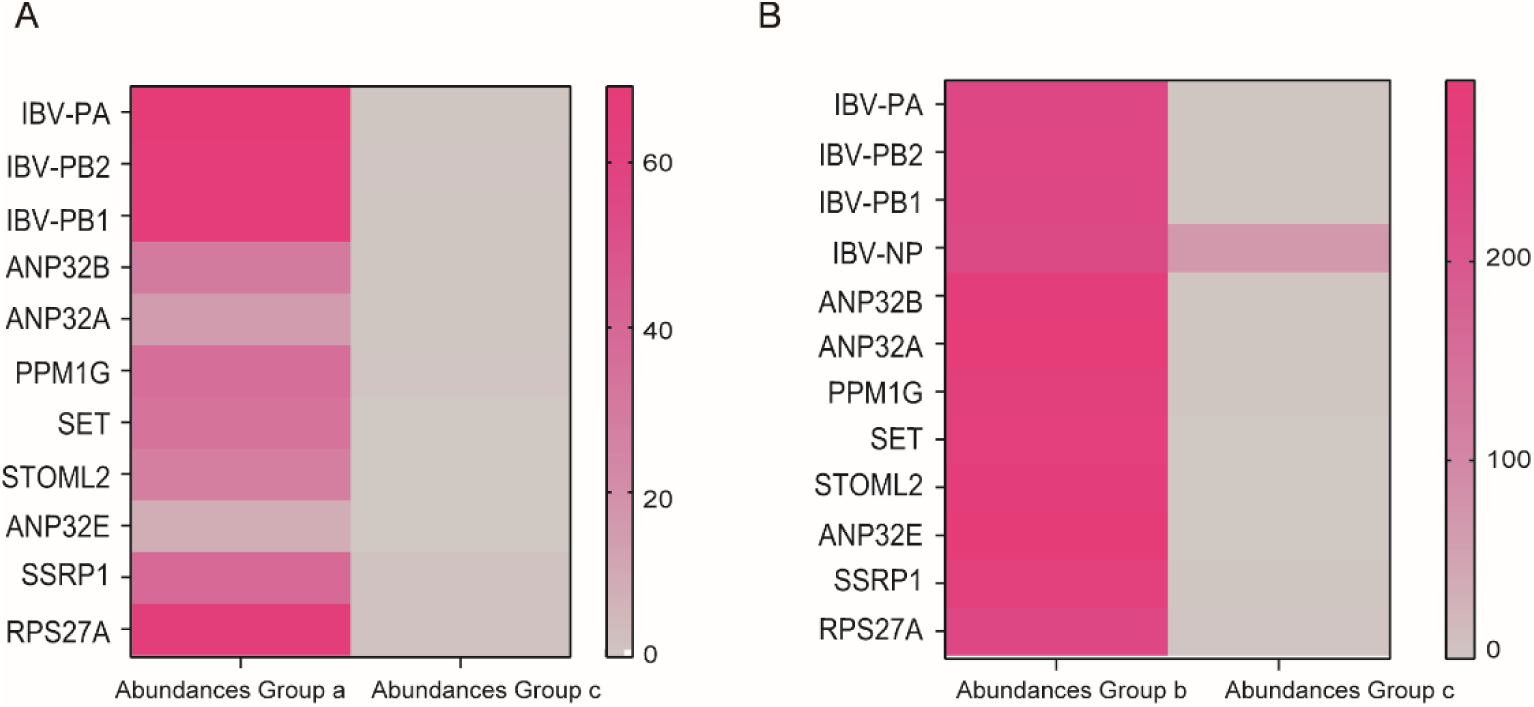
The abundance of candidate proteins in RdRP and vRNP. (A) MS analysis revealed the protein abundance of PA, PB2, PB1, ANP32B, ANP32A, PPM1G, SET, STOML2, ANP32E, SSRP1 and RPS27A in Group a versus Group c. (B) PA, PB2, PB1, NP, ANP32B, ANP32A, PPM1G, SET, STOML2, ANP32E, SSRP1 and RPS27A were quantified in Group b versus Group c by MS analysis

**Figure S2:**
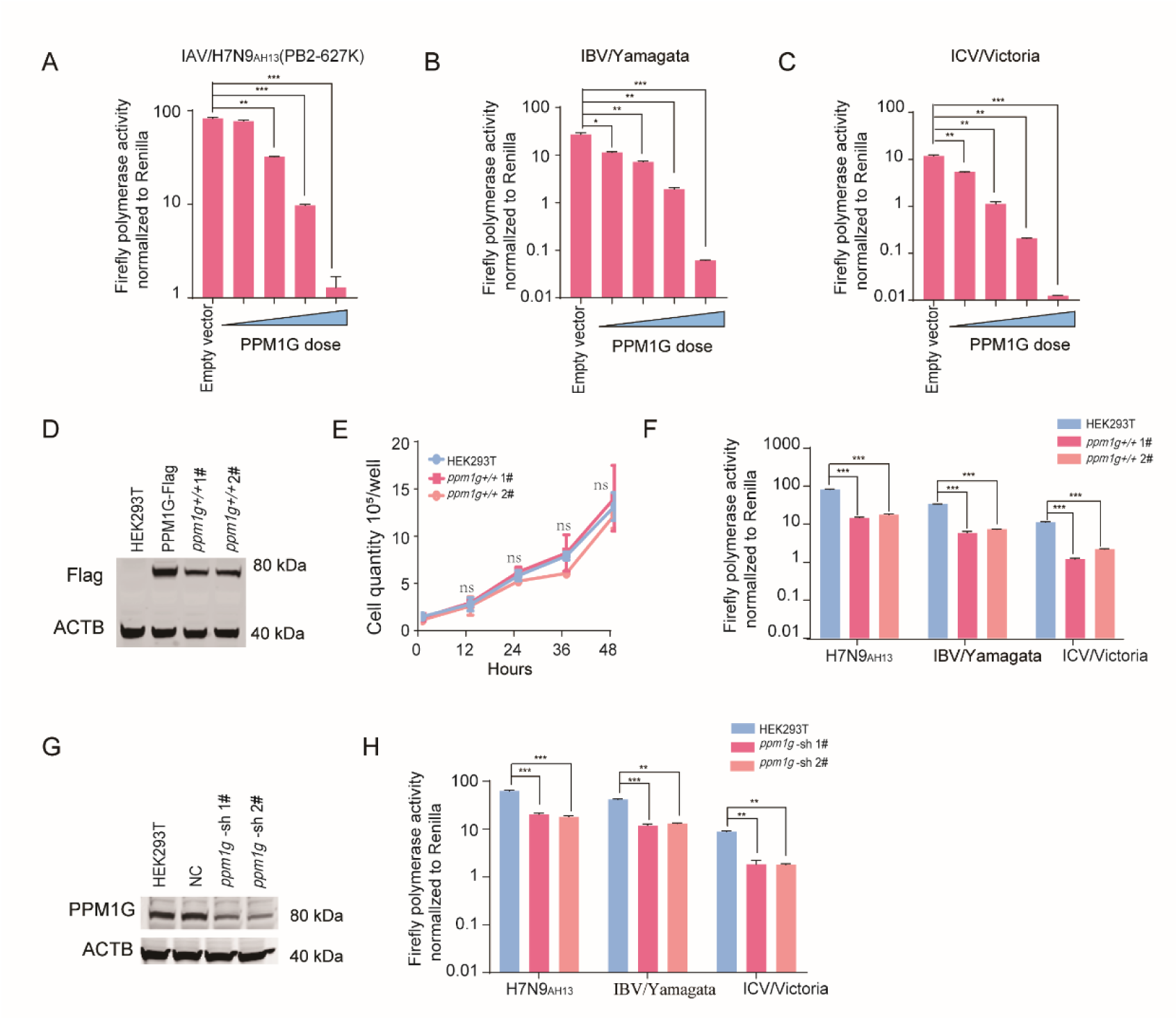
Stable overexpression and knockdown of *ppm1g* significantly inhibit influenza virus polymerase activity. A-C: Dose-dependent overexpression of PPM1G significantly inhibits the polymerase activity of H7N9_AH13_ (PB2-627K), IBV-Yamagata and ICV/Victoria. NS = not significant, \**p*<0.05, \*\**p*<0.01, *** *p* < 0.001. D-F: Detection of the expression of PPM1G protein in WT and *ppm1g*+/+ cells (HEK293T cells stably expressing *ppm1g* gene) using anti-Flag (D), and detection of their physiological status (E). Minigenome assays of the activity of polymerases from H7N9_AH13_(PB2-627K), IBV-Yamagata and ICV/Victoria in WT or *ppm1g*+/+ cells (F). Error bars represent means ± SD from n = 3 independent biological replicates; Statistical significance was determined by two-tailed unpaired t-test. NS = not significant, *** *p* < 0.001. G, H: Detection of the expression of PPM1G protein in WT or sh-*ppm1g* cells (G). Minigenome assay of activities of polymerases from H7N9_AH13_(PB2-627K), IBV-Yamagata and ICV/Victoria in WT and sh-*ppm1g* cells (H). Error bars represent means ± SD from n = 3 independent biological replicates; statistical significance was determined by two-tailed unpaired t-test. \*\**p*<0.01, *** *p* < 0.001.

**Figure S3:**
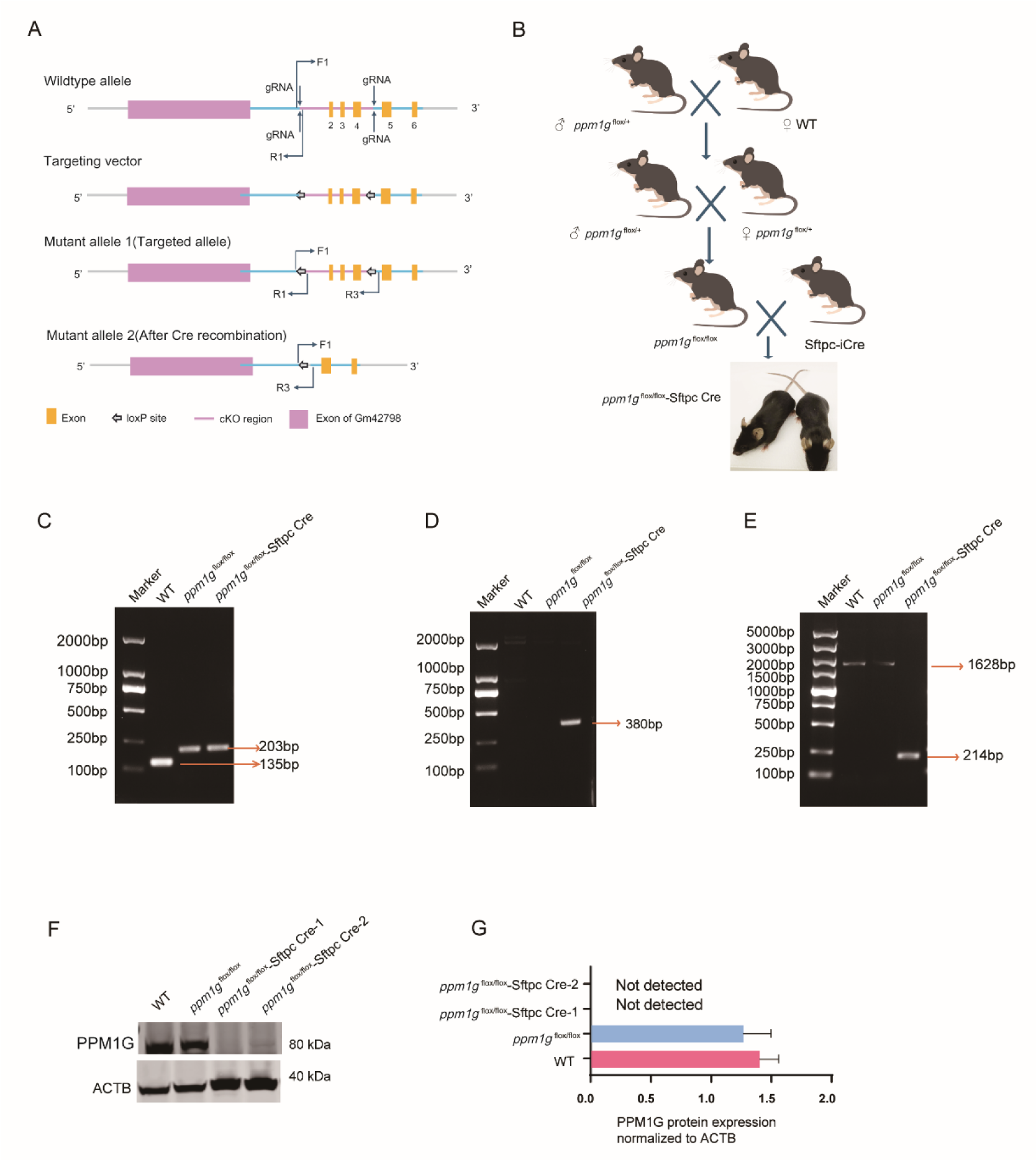
Identification of *ppm1g* ^flox/flox^-Sftpc Cre mice. Schematic diagram of targeting strategy (A) and breeding (B) for *ppm1g* ^flox/flox^-Sftpc Cre mice. C-E: Genotypes of WT and *ppm1g* ^flox/flox^-Sftpc Cre mice confirmed by PCR. (C) when using the primers 1(F1 5’-AGGAAGGGTCTTCATGTGCCATA-3’ and R1 5’- CAAATGATGGCTCAGAGTTAAGGTG-3’) for genotyping, the WT and *ppm1g*^flox/flox^ mice produced a 135 bp band, while the homozygous *ppm1g*^flox/flox^ Cre mice generated a 203 bp band (up panel). (D) PCR with primers 2 (Sftpc-cre F 5’- ATCTCTTAAAGTCAGTGGTCTCGG -3’and Sftpc-cre R 5’-CATTCAACAGACCTTGCATTCCTT-3’) showed no band in WT and *ppm1g*^flox/flox^ mice, while a 380 bp band was generated in homozygous *ppm1g*^flox/flox^ Cre mice. (E) PCR with primers 3 (F1 5’-AGGAAGGGTCTTCATGTGCCATA-3’and R3 5’-CAT-ACTTCCCCATTGTACCTCTCC-3’), the WT and *ppm1g*^flox/flox^ mice produced a 1628 bp band, the homozygous *ppm1g*^flox/flox^ Cre mice yielded a 214 bp band. F: WB detection of PPM1G protein expression in the lungs of WT, *ppm1g*^flox/flox^ and *ppm1g*^flox/flox^- Sftpc Cre mice. PPM1G protein expression was quantified from WB images using ImageJ (G).

**Figure S4:**
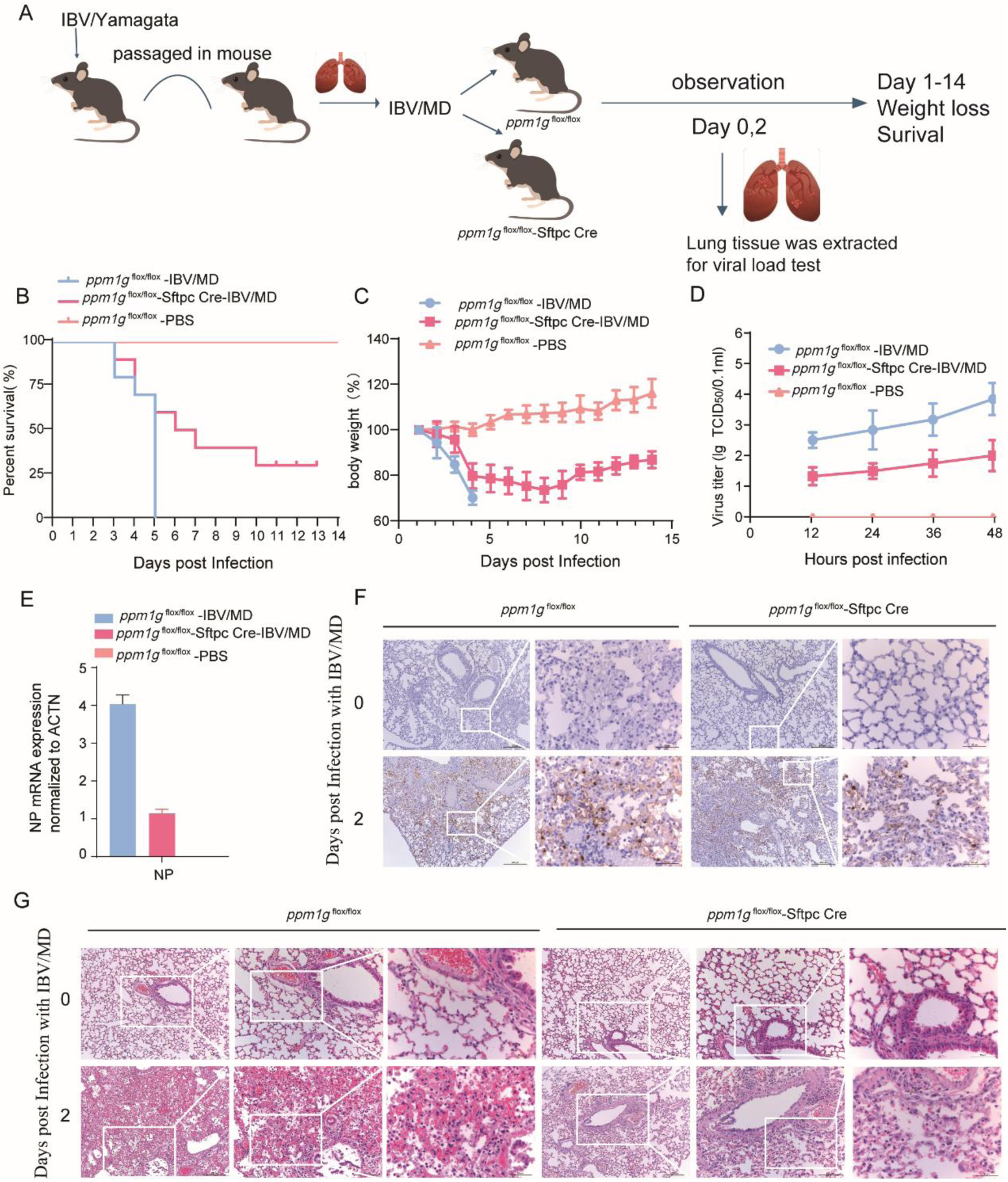
*ppm1g*-deficient mice are resistant to IBV infection. A-C: The IBV virus was passaged in mice to obtain an IBV adaptive strain (IBV-MD) that could cause death, and an infection model was established (A), the survival rate (B) and weight changes (C) of the *ppm1g* ^flox/flox^-Sftpc cre (6 weeks, n=10)and *ppm1g* ^flox/flox^ mice(6 weeks, n=10) were monitored. D: Viral titer in lung tissues from IBV-MD virus infected *ppm1g* ^flox/flox^-Sftpc cre and *ppm1g* ^flox/flox^ mice at 2 dpi as determined by TCID_50_ assays. Data are from three independent experiments (n = 3 mice per group) run in triplicate. E: RT-PCR analysis of viral NP mRNA levels in *ppm1g* ^flox/flox^-Sftpc cre and *ppm1g* ^flox/flox^ mice lung tissues. F-G: H&E staining (F) and NP immunohistochemistry (G) of lung tissues from *ppm1g* ^flox/flox^-Sftpc cre and *ppm1g* ^flox/flox^ mice infected with virus at 2 dpi; scale bars=50 or 200 μm. Representative images from 3 mice per group of three independent experiments.

**Figure S5:**
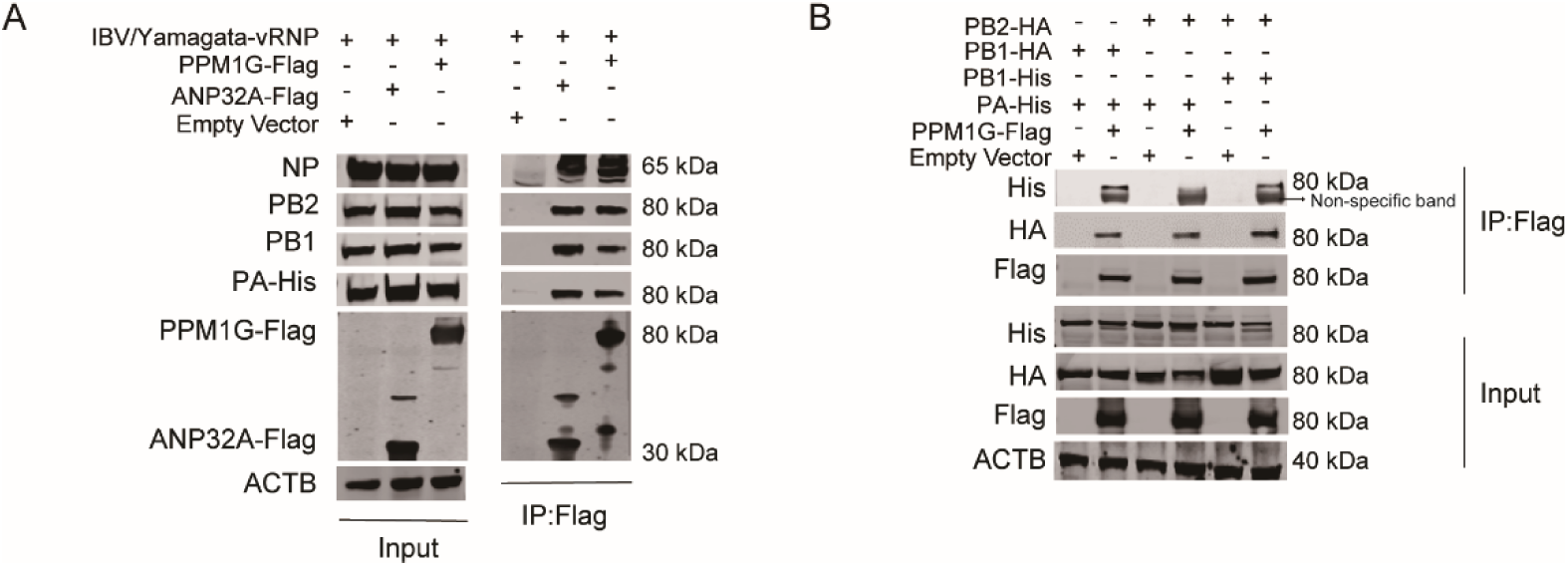
PPM1G interacted with IBV/Yamagata vRNP and PA+PB1/PB1+PB2. A: IBV/Yamagata vRNP (PA, PB1, PB2, NP and vLuc) were co-transfected into HEK293T cells with either ANP32A-Flag or PPM1G-Flag. Protein binding was detected by Co-IP, and protein expression was analyzed by Western blotting. B: IBV/Yamagata PA-His+PB1-HA, PA-His+PB2-HA and PB1-His+PB2-HA were each co-transfected into HEK293T cells with PPM1G-Flag. Protein binding was detected by Co-IP, and protein expression was analyzed by Western blotting.

**Figure S6:**
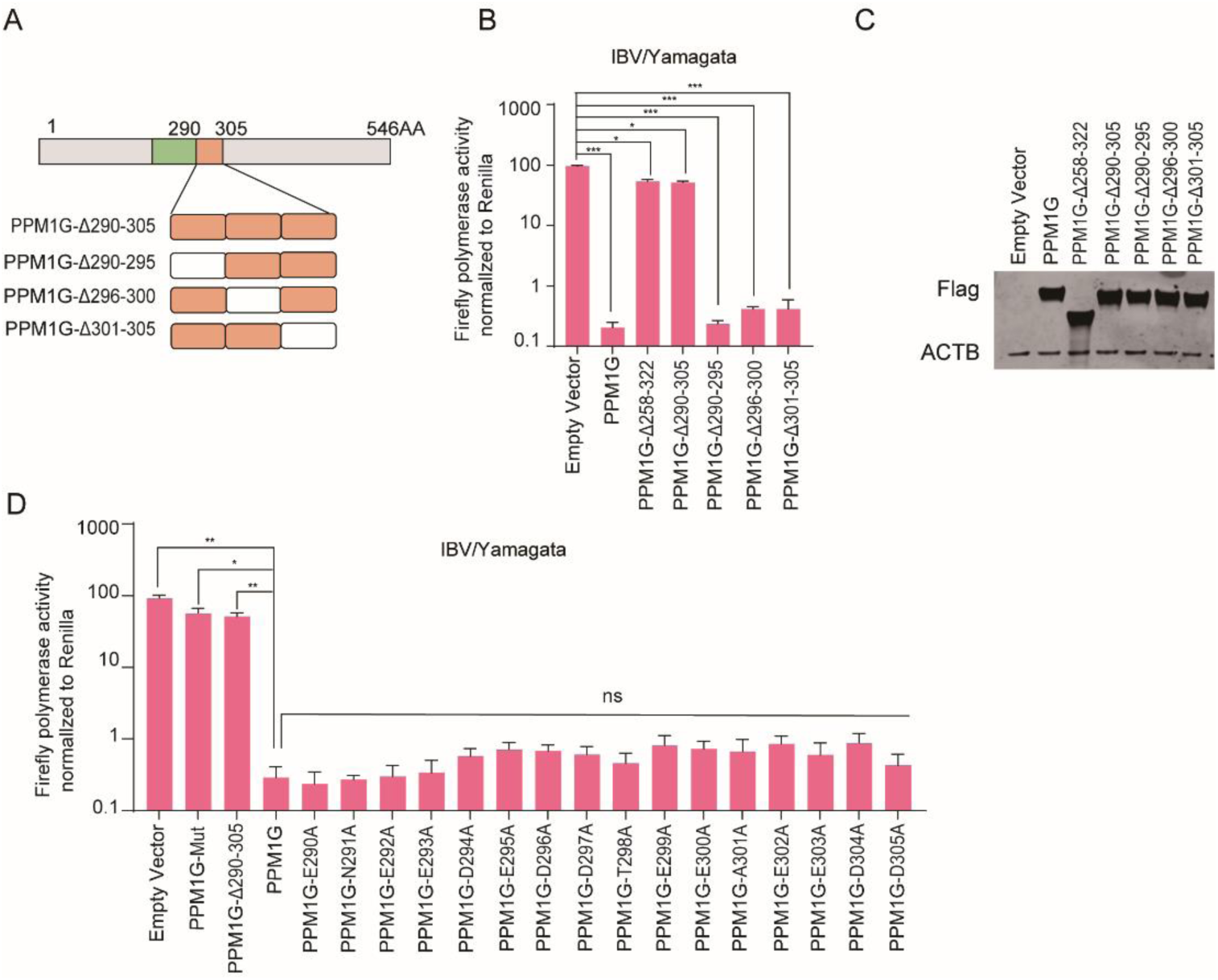
The 290-305 domain of PPM1G is the smallest domain to regulate the activity of influenza virus polymerases. A-C: Schematic diagram of the 290-305 amino acid domain of PPM1G truncated into 3 segments (A). Each of these 5 mutant proteins were transfected separately into HEK293T cells together with the IBV/Yamagata minigenome reporting system. Luciferase activity was measured 24 h after transfection (B). Analysis of expression of PPM1G mut, PPM1G-Δ-290-305 mut, PPM1G-Δ-290-295 mut, PPM1G-Δ-296-300 mut, PPM1G-Δ-301-305 mut proteins with WB (C). Error bars represent the SD of the replicates within one representative experiment. NS = not significant, \**p*<0.05, *** P < 0.001. D: There are 16 amino acid point mutations in PPM1G-290-305 mut. These 16 eukaryotic expression plasmids were co-transfected into human HEK293T cells with the IBV-Yamagata minigenome reporting system. NS = not significant, \**p*<0.05, ** *p* < 0.01.

**Figure S7:**
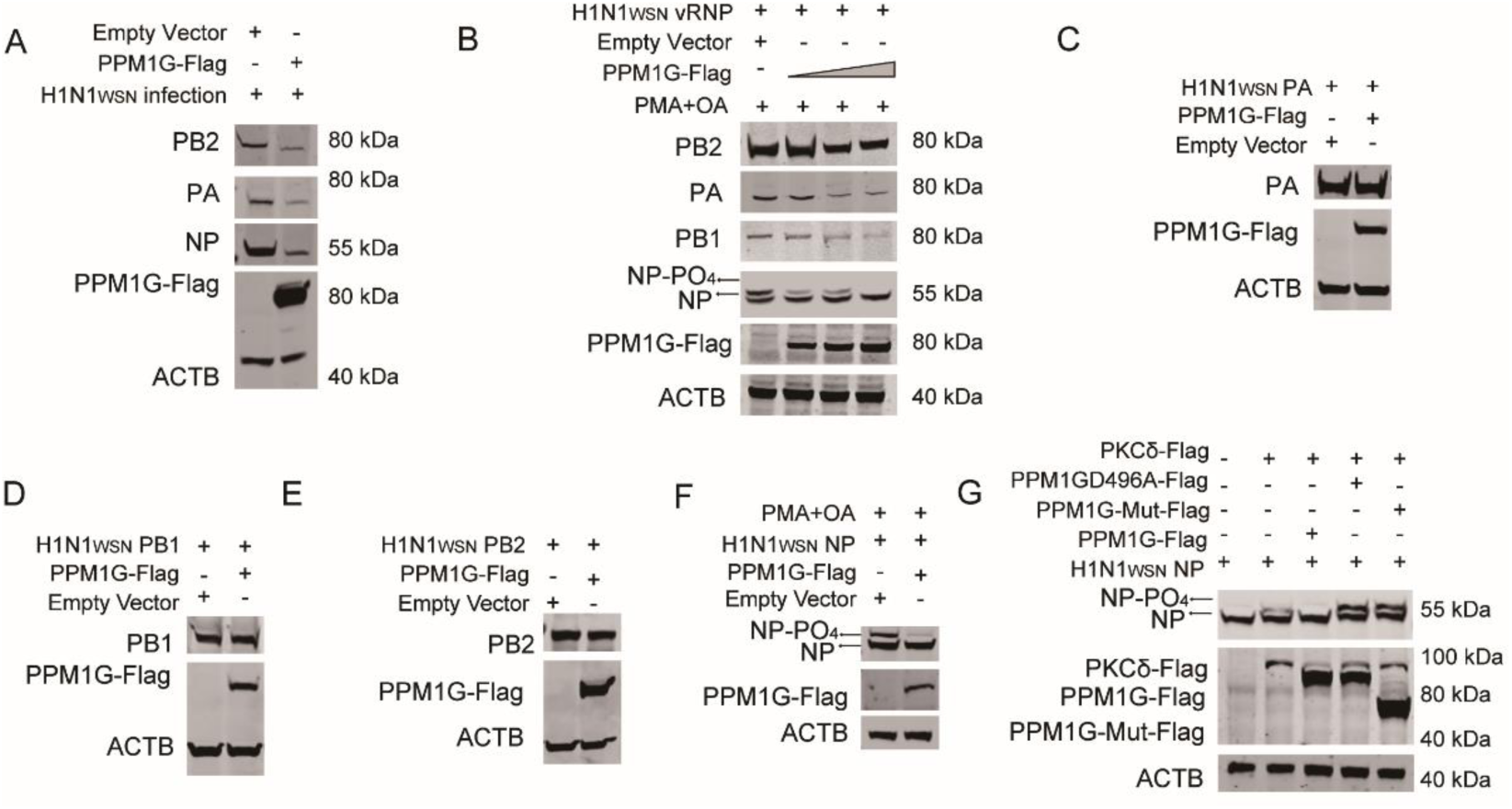
PPM1G dephosphorylates IAV-NP. A, B: A549 cells overexpressing PPM1G-Flag or empty control were infected with IAV/H1N1_WSN_ (MOI=1) (A) or were transfected with the H1N1_WSN_ vRNP (B). The vRNP protein levels in cell lysate was then determined on a Phos-tag™ SDS-PAGE. C-F: PPM1G-Flag was co-transfected with H1N1_WSN_ PA-His (C), PB1-His (D), PB2-His (E) or NP- His (F) into HEK293T cells (PMA and OA were added). Changes in the level of proteins were then assessed by Western blot using Phos-tag™ SDS-PAGE G: H1N1_WSN_ NP-His and PKCδ were co transfected into HEK293T cells with PPM1G-Flag, PPM1G-Mut-Flag or PPM1G-D496A-Flag. The change in levels of IAV-NP phosphorylation was determined using a Phos-tag™ SDS-PAGE.

**Figure S8:**
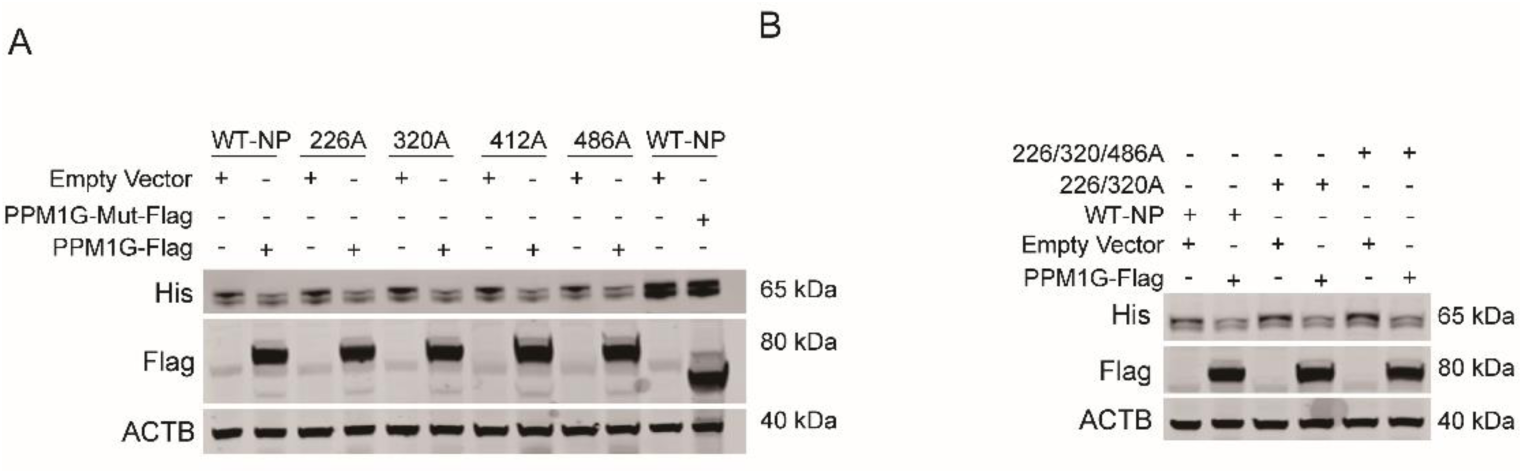
NP-S226, NP-S320, NP-S412 and NP-S486 single-point mutations and three-point joint mutations are all regulated by PPM1G. A, B: PPM1G-Flag was co-transfected into HEK293T cells together with empty vector, NP-S226A, NP-S320A, NP-S412A, NP-S486A, NP-S226A/320A or NP-S226A/S320A/S486A. NP phosphorylation form was assessed using WB.

